# B cell/adipocyte crosstalk drives pro-fibrotic macrophage migration and polarisation in Crohn’s disease-associated, inflamed adipose tissue

**DOI:** 10.1101/2025.07.11.664132

**Authors:** Marie Christine Goess, Julia M. Heinze, Julian Hofmann, Annalisa Buck, Iana Gadjalova, Marie-Christin Weber, Sandra Ring, Joerg M. Buescher, Matteo Villa, Philipp-Alexander Neumann, Jürgen Ruland, Selina J. Keppler

## Abstract

During Crohn’s disease (CD), hyperplasia of the mesenteric white adipose tissue (WAT), called creeping fat (CrF), is associated with the accumulation of immune cells. We here investigated whether B cell infiltration and their interaction with adipocytes influence inflammation and fibrosis of CD-associated adipose tissue. Analysing CrF of CD patients as well as WAT from a mouse model of intestinal inflammation, we found an accumulation of B cells and pro-fibrotic, M2-like macrophages. Depleting B cells through anti-CD20 antibody treatment diminished M2-like macrophage accumulation in inflamed mouse WAT. Mechanistically, we found that the B cell/adipocyte co-cultures led to elevated secretion of monocyte chemoattractant protein-1 (MCP-1) by primary adipocytes, in part due to TNF-a secretion by activated B cells. Adipocyte-derived MCP-1 resulted in enhanced macrophage migration. Furthermore, the supernatants of the B cell/adipocyte co-culture promoted pro-fibrotic, M2-like macrophage polarization *in vitro*, correlating with elevated levels of lactate. Single-cell RNA sequencing of human CrF and mouse WAT supported our *in vitro* findings suggesting that the B cell/adipocyte crosstalk supports a lactate-rich, inflammatory adipose tissue niche. Taken together, our results provide evidence for a crucial role of the B cell/adipocyte crosstalk to CD-associated adipose tissue inflammation.

**Graphical abstract:** 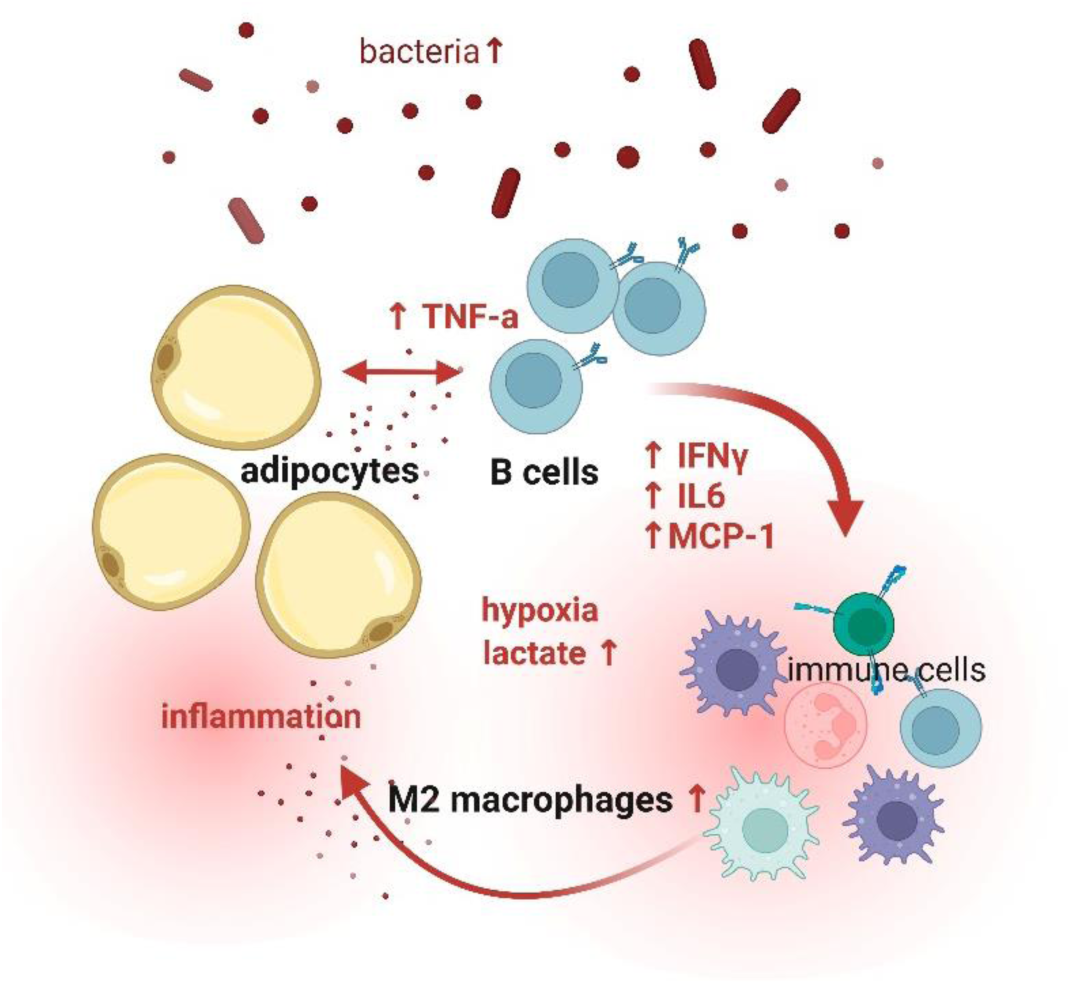

**One Sentence Summary:**

- B cells and pro-fibrotic macrophages accumulate in IBD-associated, inflamed adipose tissue
- B cell depletion in a mouse model reduces pro-fibrotic macrophage accumulation in IBD-associated, inflamed adipose tissue
- B cell/adipocyte crosstalk promotes B cell activation and MCP-1 secretion by adipocytes
- Adipocyte-derived MCP-1 promotes migration of macrophages
- B cell/adipocyte crosstalk provides a lactate-rich, inflammatory environment which likely drives pro-fibrotic macrophage polarisation

## Introduction

Inflammatory bowel disease (IBD), including Crohn’s Disease (CD) and Ulcerative Colitis (UC), is a group of immune-mediated disorders of the intestine. During IBD, a barrier breach of the intestinal epithelium results in microbial translocation into the surrounding tissue, including the mesenteric white adipose tissue (MAT) (1), resulting in inflammation. During CD, the inflamed MAT displays adipocyte hyperplasia (2) eventually leading to fat-wrapping around the intestine (so called “creeping fat” (CrF) (3)). CrF is an extra-intestinal inflammation of CD, which is thought to restrict bacterial dissemination and the spread of inflammation, but also correlates with complications, such as intestinal fibrosis and stricture formation (4). Adipocytes in CrF are smaller (5, 6) and seem to be metabolically dysfunctional shifting from fatty acid synthesis to glycolysis and showing signs of hypoxia (7). CrF of CD patients has been demonstrated to favour the differentiation and accumulation of pro-fibrotic, M2-like, CD206 expressing macrophages (1, 8). The presence of bacteria in the MAT can be sensed by those adipose-tissue macrophages (1, 8), but also by adipocytes themselves (9) resulting in the production of pro-inflammatory cytokines such as Interleukin-6 (IL-6), Tumour necrosis factor alpha (TNF-α), Interleukin-1 (IL-1), and monocyte chemoattractant protein (MCP-1, also known as C-C chemokine ligand 2 (CCL2)) (7, 9). This local release of pro-inflammatory cytokines might lead to the recruitment and activation of immune cells. Studies of CD patients report immune cell aggregates resembling tertiary lymphoid organs around lymphatic vessel of the MAT, adjacent to areas of intestinal inflammation (8, 10, 11). A recent transcriptomics study of CrF identifies markers of B cells, T cells as well as plasma cells associated with CD (1, 12). However, how the reciprocal interplay between inflamed adipocytes and infiltrating lymphocytes affects MAT inflammation remains incompletely understood.

B cells might influence MAT inflammation through antibody and cytokine production or interaction with T cells. During obesity-induced or aging-associated adipose tissue inflammation, white adipose tissue (WAT) contains antibody secreting and pro-inflammatory cytokine producing B cells (13-16). While the phenotype and function of infiltrating immune cells during diet-induced or age-associated obesity has been studied extensively, little is known about the phenotype and function of WAT infiltrating B cells during inflammatory, non-obese conditions, such as during IBD.

Here, we investigated how the reciprocal interplay between inflamed adipocytes and B cells influences IBD-associated adipose tissue inflammation. We established a spheroid culture system to study adipocyte inflammation in vitro. We found that CrF of CD patients display an inflammatory memory, which can be mimicked by LPS-stimulation of human and mouse primary adipocytes *in vitro*. Furthermore, our results suggest that WAT infiltrating B cells enhanced MCP-1 secretion by adipocytes resulting in increased macrophage migration. In addition, B cell/adipocyte crosstalk provided a lactate-rich, inflammatory environment which supported pro-fibrotic M2-like macrophage polarization. Concomitantly, depletion of B cells in our mouse model of intestinal inflammation reduced the accumulation of the M2-like macrophage population in inflamed WAT. Taken together, our results provide evidence for a crucial role of the B cell/adipocyte crosstalk to IBD-associated adipose tissue inflammation.

## Results

### B cells accumulate in human CrF and mouse inflamed WAT during intestinal inflammation

To characterize the composition of immune cells in adipose tissue in CD patients on a cellular level, we analysed biopsies from CrF around inflamed and stenotic intestinal segments and compared them to uninvolved MAT from the resection margin from the same patient (Fig. 1A, patients’ characteristics summarised in Table 1 and Supplemental data Table 1). Using multi-colour flow-cytometry, we observed a significantly increased number of CD45-expressing leukocytes in CrF tissue compared to MAT (Suppl. Fig. 1A +B), with an overabundant population of CD19^+^ B cells (Fig. 1B+C), IgA^+^, CD19^low^ plasma cells (Suppl. Fig.1C+D) and IgA^+^, CD19^high^ plasmablasts (Suppl. Fig.1C+E). Our analysis also revealed a modestly increased presence of CD3 T cells (Suppl. Fig. 1F+G), and a variable amount of macrophages (Suppl. Fig.1H+I) in CrF relative to MAT. Thus, we found accumulation of different immune cells, especially B cells and plasma cells, in CrF compared to MAT in CD patients.

**Figure 1:**
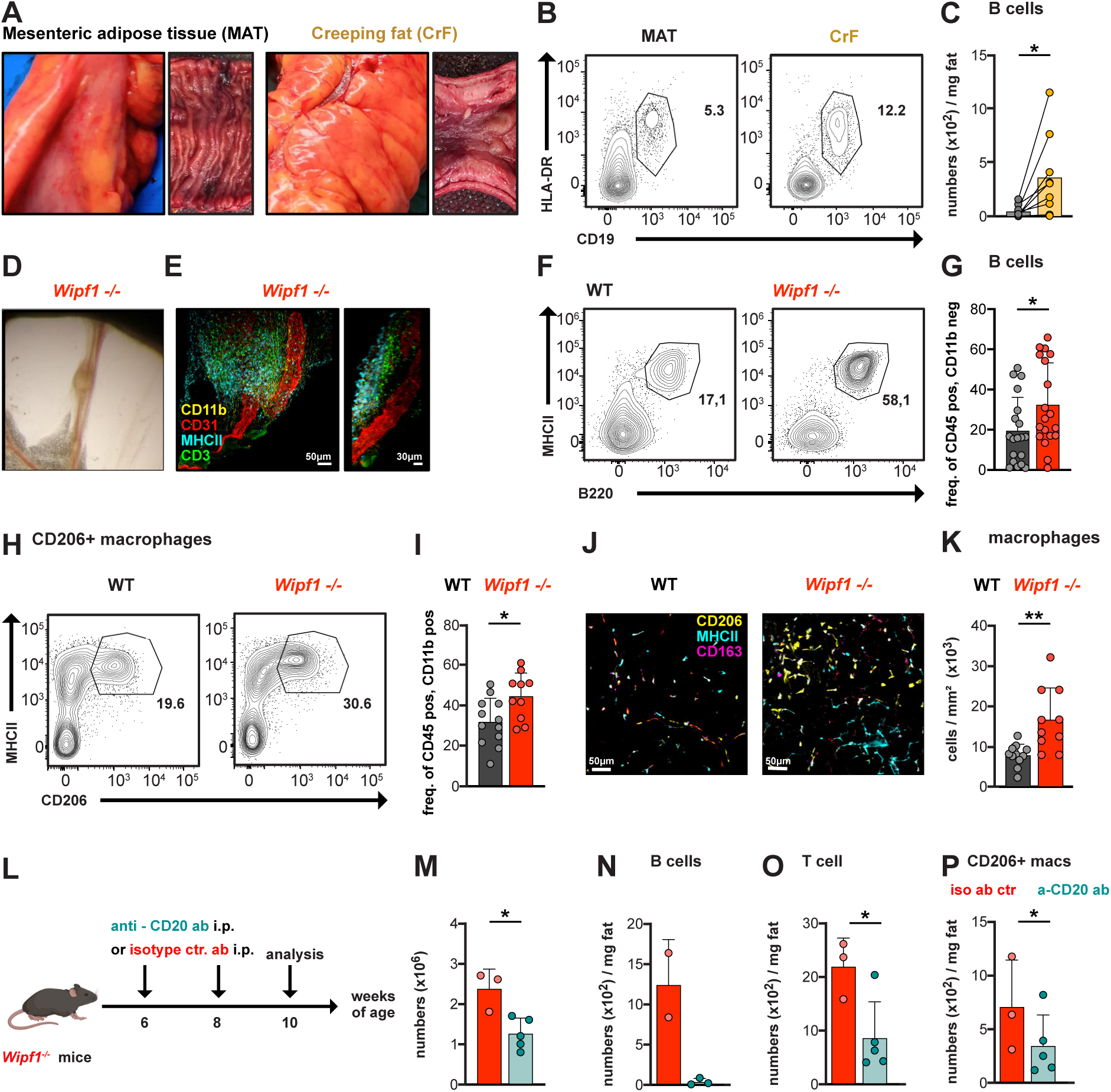
B cell depletion reduces pro-fibrotic, M2-like macrophage accumulation in WAT in a mouse model of IBD. (A) Representative images of MAT of a CD patient. Left side: Uninvolved intestinal segment (MAT, external and mucosal perspectives). Right side: macroscopic characteristics of a small bowel segment with attached CrF and a stricture visualized from the mucosal aspect. (B) Representative flow plots of human CD45^+^, CD11b^−^, CD19^+^, HLA-DR^+^ B cells, quantified as cell counts per mg fat isolated from MAT and CrF (C). *n≥7 patients.*(D+E) Micrograph and immunofluorescent image (E) of a lymphoid structure in MAT of *Wipf1^−/−^*mice. (F) Representative flow plots of CD45^+^, CD11b^−^, B220^+^, MHC-II^+^ B cells isolated from visceral WAT from WT and *Wipf1^−/−^* mice quantified in (G) (frequencies, mean ± SD). (H) Representative flow plots of CD45^+^, CD11b^+^, CD206^+^, MHC-II^+^ M2 macrophages quantified in (I) (frequencies, mean ± SD) (F-I) *n≥10 mice per genotype, pooled from multiple independent experiments.* (J) Confocal microscopy of cleared adipose tissue of WT and *Wipf1^−/−^* mice showing CD206^+^ (yellow), CD163^+^ (magenta), and MHC-II^+^ (cyan) monocytes/macrophages quantified in (K) (mean ± SD) n=6 mice/ genotype; 1-2 3D images/mouse. (L) Experimental design of B cell depletion in *Wipf1* ^−/−^ mice using anti-CD20 or isotype control antibodies. (M) Total cell numbers isolated from WAT. Total number/mg WAT of (N) CD19⁺,MHC-II⁺ B cells, (O) CD3⁺,CD4⁺ T cells, and (P) CD206⁺,MHC-II⁺ macrophages isolated from WAT of treated *Wipf1* ^−/−^ and control mice. n ≥ 3 per group, pooled from two independent experiments. *Statistical significance was calculated(C, M-P) paired and (G,I,K) unpaired student’s t test. *p < 0.05; **p < 0.01; ***p < 0.001; ****p < 0.0001. Illustration 1L was created with BioRender.com*.

**Table 1.** Patients’ characteristics – summary.

| **Patients` Characteristics** |  | **n= 17** |
| --- | --- | --- |
| Age at time of surgery (years) Median [IQR] |  | **50 [21]** |
| Age at diagnosis (years) Median [IQR] |  | **24 [5]** |
| Gender n [%] | **Male** | **10 [59]** |
|  | **Female** | **7 [41]** |
| Disease behavior n [%]<br>Montreal classification | **B1: Nonstricturing** | **0 [0]** |
|  | **B2: Stricturing** | **17 [100]** |
|  | **B3: Penetrating** | **0 [0]** |
| Localization [%] | **Terminal ileum** | **15 [88]** |
|  | **Colon** | **2 [12]** |
| Medication at the timepoint of surgery n [%] | **Adalimumab** | **1 [6]** |
|  | **Infliximab** | **2 [12]** |
|  | **Ustekinumab** | **6 [35]** |
|  | **Glucocorticoids** | **6 [35]** |
|  | **None** | **2 [12]** |
| BMI Median [IQR] |  | **23 [2]** |
| Smoking status n [%] | **Smoker** | **4 [24]** |
|  | **Non-smoker** | **13 [76]** |

We have previously described mice lacking the Wiskott-Aldrich Syndrome protein interacting protein (*Wipf1^−/−^* mice), a genetically engineered mouse that develops colitis, as model for human IBD (17). Examining the visceral WAT of *Wipf1^−/−^* mice (Suppl. Fig. 1J), we observed the formation of lymphoid structures along the blood vessels in the IBD-affected mesentery (Fig. 1D), concomitant with the accumulation of immune cells (Fig. 1E). Using flow cytometry, we found a significant increase in B cells (Fig. 1F+G, Suppl. Fig. 1K) and a notable, though not statistically significant, rise in T cells (Suppl. Fig. 1L+M) isolated from inflamed WAT compared to homeostatic tissue, similar to our findings in human biopsies. Concomitantly, we found a substantial population of CD206-expressing, M2-like macrophages (Fig. 1H+I) in inflamed WAT compared to controls, which we confirmed using tissue clearing and volumetric imaging (Fig. 1J+K)(18). Together, the infiltration of inflammatory immune cells in IBD-associated WAT, similar to what we observed in CrF of CD patients, indicated a prominent adipose tissue inflammation in the *Wipf1^−/−^* mouse model of intestinal inflammation.

### B cell depletion reduces pro-fibrotic, M2-like macrophage accumulation in WAT in Wipf1^−/−^ mice

Having observed an enhanced presence of B cells in human CrF and mouse WAT, we went on to directly assess the role of B cells during WAT inflammation in *Wipf1^−/−^* mice. For this, we depleted B cells of *Wipf1* ^−/−^ mice at 6 and 8 weeks of age using an anti-CD20 antibody or an isotype control antibody (Fig. 1L). WAT was analysed at 10 weeks of age. B cell depletion did not affect the weight of WAT between anti-CD20 or isotype control treated groups (Suppl. Fig. 1N). Nevertheless, total CD45+ leukocyte counts were significantly reduced in WAT of B cell-depleted *Wipf1* ^−/−^ mice (Fig. 1M), concomitant with efficient B cells depletion within WAT (Fig. 1N, Suppl. Fig. 1O). Notably, B cell depletion also resulted in a 50% reduction of CD3 T cell numbers (Fig. 1O, Suppl. Fig. 1P). B cell depletion compared to isotype control treatment did not significantly affect the frequencies and cell number of CD11b cells isolated from WAT (Suppl. Fig. 1Q+R), but led to an about 50% reduction of cell numbers of CD206 macrophages (Fig. 1P, Suppl. Fig. 1S). Thus, the depletion of B cells in our mouse model of intestinal inflammation led to a reduced presence of T cells and M2-like macrophages in IBD-associated WAT. We concluded that B cells might play a prominent role in the recruitment of immune cell populations and the accumulation of a M2-like macrophage population during an early stage of WAT inflammation.

### WAT-derived spheroids model IBD-related adipose tissue inflammation in vitro

Adipose tissue is composed of mature adipocytes and the stromal vascular fraction (SVF). The SVF contains immune cells and a heterogeneous population of mesenchymal stem cells, including committed pre-adipocytes that can differentiate into adipocytes (19). To investigate human adipose tissue inflammation, as well as the interaction with immune cells, in more detail *in vitro,* we used a pre-adipocyte spheroid culture system. For this, we isolated the SVF of human MAT or CrF (patients’ characteristics summarised in Table 1, Supplemental data Table 2), expanded the SVF into pre-adipocytes, cultured them as self-organizing 3D spheroids followed by differentiation into adipocytes (Fig. 2A). Lipid uptake was verified using BODIPY staining (Fig. 2B). Secreted cytokines in the culture supernatant were measured using a multiplex assay. We detected a significantly higher concentration of the pro-inflammatory cytokine IFN-γ, consistent but not significantly elevated concentrations of IL-6 and of the monocyte/macrophage chemoattractant protein (MCP-1), and very little TNF-α or IL-10 in the culture supernatants of CrF-derived adipocyte spheroids compared to MAT-derived adipocyte spheroids (Fig. 2C).

**Figure 2:**
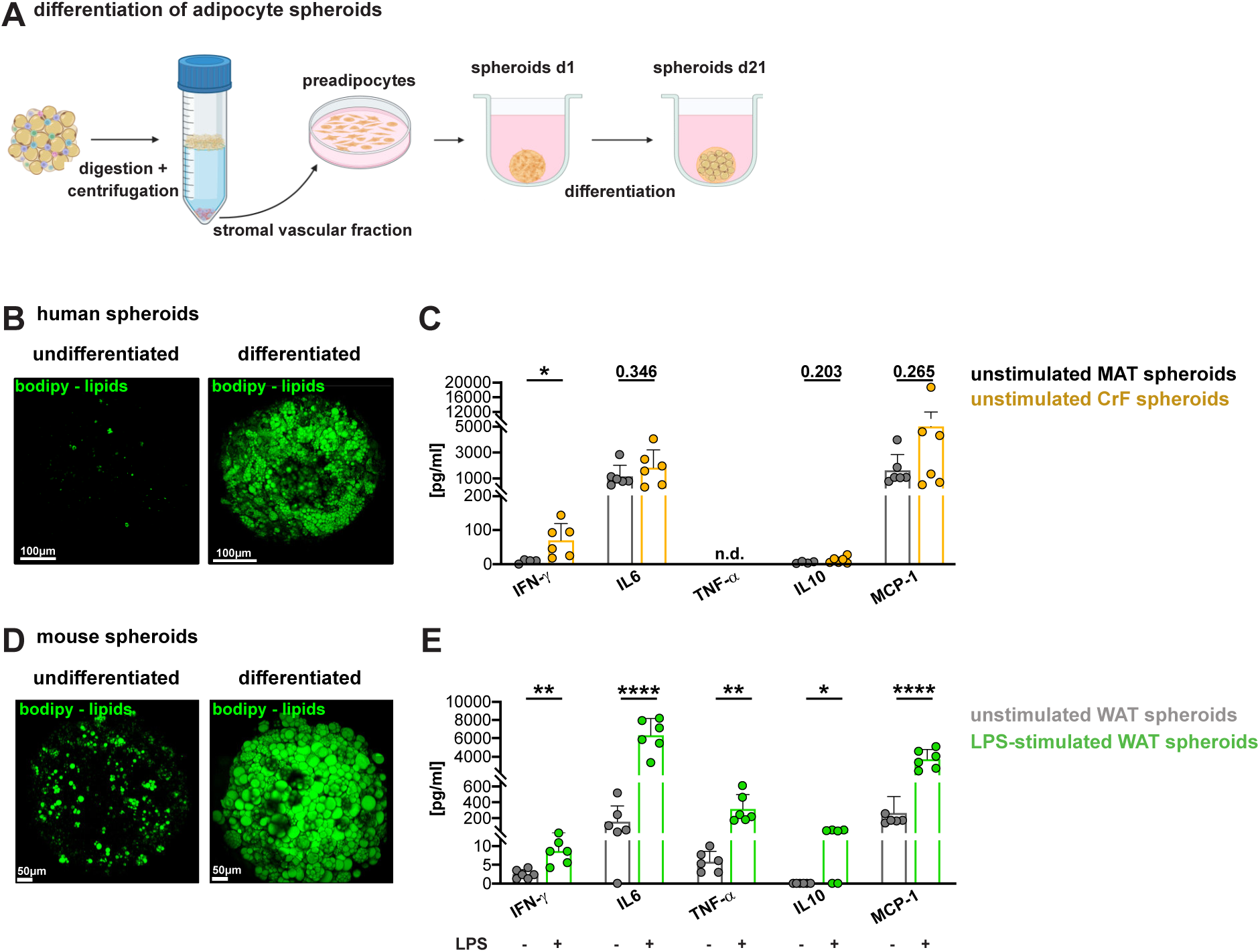
WAT-derived spheroids model IBD-related adipose tissue inflammation in vitro. (A) Experimental setup for adipose tissue-derived spheroid cultures. (B) Representative immunofluorescence images of human spheroids at day 1 (undifferentiated spheroids, left) and day 21 (differentiated spheroids, right) stained with BODIPY (green) for lipid visualization. (C) Cytokine secretion profile in the supernatants of differentiated MAT- or CrF-derived adipocyte spheroids as measured using a multiplex assay (mean ± SD). For comparison purposes, cytokines of supernatants of unstimulated MAT-derived spheroids plotted are the same in Fig. 2C and Suppl. Fig. 2B. *Spheroids were generated from n≥5 patients, with 2 distinct tissue groups (MAT, CrF) per patient. Statistical significance was calculated using unpaired student’s t test.* (D) Representative immunofluorescence images of murine adipocyte spheroids at day 1 (undifferentiated spheroids, left) day 21 (differentiated spheroids, right) stained with BODIPY (green) for lipid visualization. (E) Cytokine secretion profile in the supernatants of differentiated unstimulated or LPS-stimulated WAT-derived adipocyte spheroids as measured using a multiplex assay (mean ± SD). *Spheroids were generated from n≥5 WT mice, pooled from 3 independent experiments. Statistical significance was calculated using unpaired student’s t test. *p < 0.05;**p < 0.01; ***p < 0.001; ****p < 0.0001. Illustration 2A was created with BioRender.com*.

The formation of CrF has been suggested to be a response mechanism against bacterial translocation from a leaky intestinal barrier (1). Correspondingly, we also found an increased abundance of bacterial DNA in IBD-associated WAT of *Wipf1^−/−^* mice compared to heterozygous littermate controls (Suppl. Fig. 2A). Consequently, to mimic bacterial infiltration, we next stimulated human MAT-derived adipocyte spheroids with lipopolysaccharide (LPS), a bacterial cell wall component (Suppl. Fig. 2B). We detected a about 10x higher concentration of IL-6 (from 1000 pg/ml to 10000 pg/ml) and MCP-1 (from 1500 pg/ml to 15000 pg/ml), and enhanced concentrations of IFN-γ, but very little TNF-α or IL-10 in the culture supernatants of LPS-stimulated compared to unstimulated human MAT-derived spheroids (Suppl. Fig. 2B). Consequently, LPS stimulation of MAT-derived adipocyte spheroids closely mimicked the cytokine secretion profile of unstimulated CrF-derived adipocyte spheroids (compare Suppl. Fig. 2B to Fig. 2C).

Due to the limited availability of human samples and in order to model human WAT inflammation in vitro, we went on to investigate murine adipocyte inflammation. First, we isolated mature adipocytes from WT mice, cultured them in a transwell culture system and stimulated them with LPS (Suppl. Fig. 2C-E). We found significantly increased levels of IL-6, TNF-α and MCP-1 secreted into the supernatant of mature adipocytes cultures in the presence of LPS (Suppl. Fig. 2E). In parallel, we isolated pre-adipocytes from WAT of WT mice (Fig. 2A), differentiated them into 3D spheroids and subsequently stimulated them with LPS (Fig. 2D). Similar to human adipocyte spheroids, LPS-stimulation of mouse WAT-derived adipocyte spheroids led to a more than 10x higher concentration of IL-6 (from 200 pg/ml to 6000 pg/ml) and MCP-1 (from 200 pg/ml to 4000 pg/ml), and enhanced concentrations of IFN-γ, TNF-α and IL-10 in the culture supernatants of LPS-stimulated compared to unstimulated mouse adipocyte spheroids (Fig. 2E). Of note, the cytokine secretion profile of mouse adipocyte spheroids after LPS stimulation (Fig. 2E) was very similar to what we have observed after LPS stimulation of human MAT-derived adipocyte spheroid cultures (Suppl. Fig. 2B); or even unstimulated CrF-derived adipocyte spheroids (Fig. 2C). Thus, murine adipocytes closely mimic human adipocyte inflammation after LPS stimulation *in vitro*.

In summary, we found that adipocyte spheroids from CrF of CD patients readily secreted enhanced levels of pro-inflammatory cytokines without additional LPS stimulation, despite the prolonged culture conditions. Furthermore, LPS stimulation of MAT-derived adipocyte spheroids mimicked the cytokine secretion profile of unstimulated CrF-derived adipocytes spheroids in our in vitro cultures. In addition, murine adipocytes closely mimic human adipocyte inflammation after LPS stimulation *in vitro*. Together, we here describe an *in vitro* culture system of differentiated adipocyte spheroids to study human, or mouse, adipocyte inflammation.

### B cell/adipocyte crosstalk promotes B cell activation and adipocyte cytokine secretion

Next, we used our in vitro adipocyte culture systems to investigate the B cell/adipocyte crosstalk. For this, we co-cultured naïve splenic *Wipf1^−/−^* B cells together with primary mature adipocytes isolated from visceral, non-inflamed WAT of WT mice for 4 days using a transwell assay and stimulated B cells and adipocytes using LPS (Suppl. Fig. 3A). We found that co-culturing B cells with adipocytes led to an enhanced survival of B cells (Suppl. Fig. 3B) and an improved differentiation into class-switched IgG1 plasmablasts (Fig. 3A+B). In addition, B cells exhibited increased surface expression of the activation marker and co-stimulatory molecule CD86 (Fig. 3C+D) when co-cultured with adipocytes, compared to LPS-stimulated B cells alone. Analysing the supernatants at day 4 of the co-culture revealed significantly higher levels of immunoglobulins IgG1, IgG2a, and IgA, but not IgG2b or IgM compared to B cells stimulated with LPS alone (Suppl. Fig. 3C). Next, we determined the concentrations of pro-inflammatory cytokines secreted by both, B cells and adipocytes into the co-culture supernatants compared to the supernatant of LPS-stimulated B cells alone. Concentrations of IL-10 and TNF-α were comparable in supernatants of both cultures (Fig. 3E). However, the concentration of IL-6 in the co-culture supernatants were increased compared to LPS-stimulated B cell alone (Fig. 3E) which is likely in parts due to adipocytes secreting IL-6 after LPS stimulation (Fig. 2E, Suppl. Fig. 2E). Interestingly, we did not detect MCP-1 in the supernatant of LPS-stimultated B cells alone (Fig. 3E+F). However, when co-cultured with adipocytes, we found consistently higher amounts of MCP-1 in the culture supernatants compared to LPS-stimulated adipocytes cultured alone (Fig. 3F). These results suggest that B cells might promote MCP-1 secretion by adipocytes in a non-contact dependent manner. To determine whether specific cytokines released by LPS-stimulated *Wipf1^−/−^*B cells might influence MCP-1 secretion by adipocytes, we used monoclonal antibodies to selectively block IL-6, IL-10, or TNF-α in co-culture experiments and compared MCP-1 secretion in each condition to that observed in co-cultures treated with an isotype control antibody. We found that only anti-TNF-α antibodies significantly reduced MCP-1 secretion by adipocytes by half when co-cultured with B cells (Fig. 3G).

**Figure 3:**
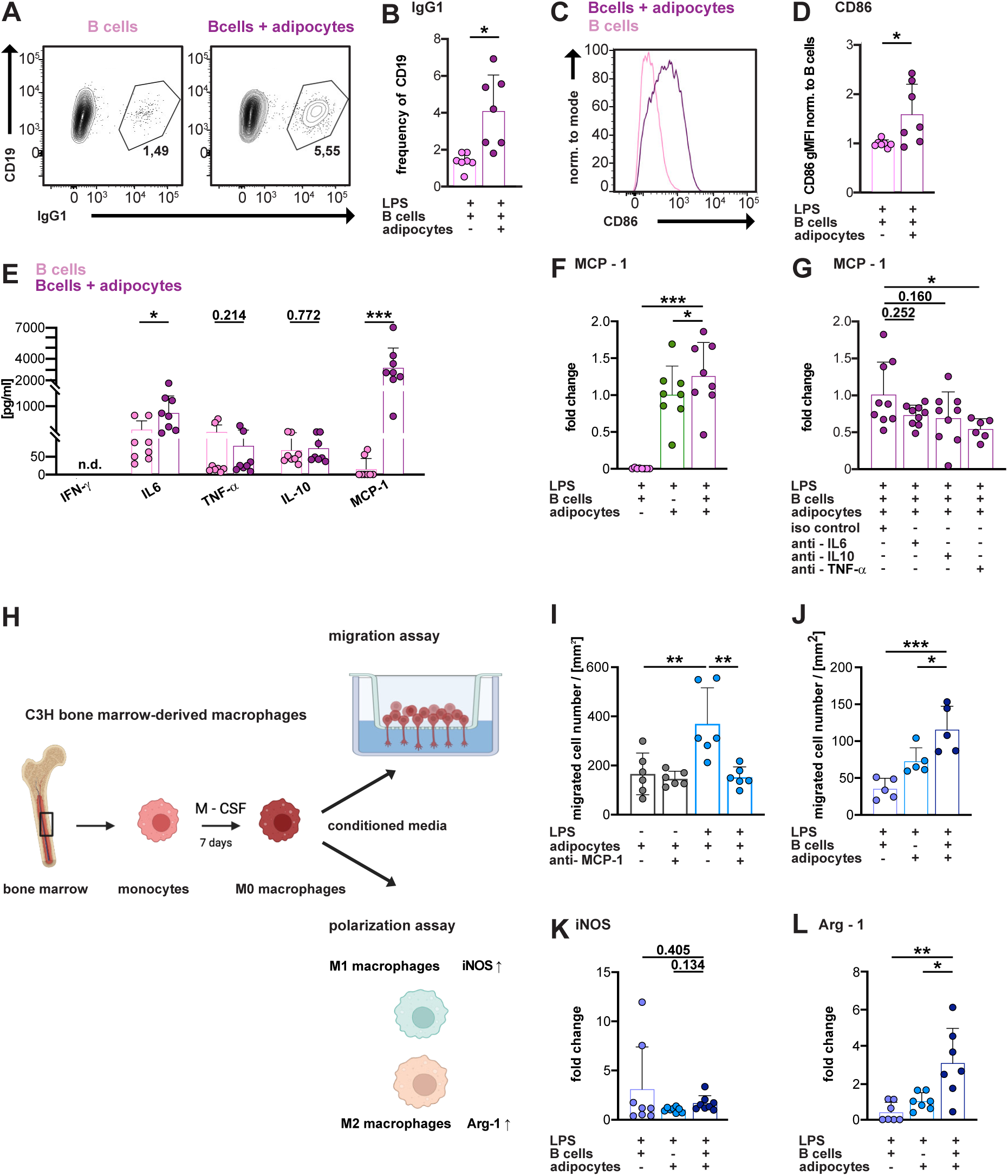
B cell/adipocyte crosstalk promotes B cell activation and adipocyte cytokine secretion. (A-D) Flow cytometric analysis of *Wipf1^−/−^* B cells cultured with or without adipocytes in the presence of LPS. (A+B) Frequencies of surface IgG1 expression. (C+D) CD86 expression on B cells (D) quantified as geometric mean fluorescence intensity (gMFI) normalized to expression on stimulated B cells. (E-G) Cytokine levels as indicated measured in the supernatants of B cell or B cell/adipocyte co-culture. (F+G) MCP-1 concentrations (F) normalised to concentration in the supernatant of LPS-stimulated adipocytes (mean fold change ± SD), or (G) in the presence of specified cytokine blocking antibodies normalized to concentrations in the the presence of the isotype control (mean fold change ± SD). *Data is shown as (B,D-E) mean ± SD. Original measurements of cytokines were in [pg/ml]. n≥7 mice pooled from 3-4 independent experiments.* (H) Experimental design of BMDM cultures, migration and polarization assays. (I+J) Quantification of macrophage migration (I) in the presence of anti-MCP-1 blocking antibody and (J) conditioned media from stimulated B cells, adipocytes, and B cell/adipocyte co-cultures. *Cells were counted in three random microscope fields per sample. Data represent mean ± SD from five mice per condition.* (K+L) Relativ mRNA expression levels (mean fold change ± SD) of (K) iNOS and (L) Arginase-1 24-hours after exposure to conditioned media as indicated, normalized to HPRT and mRNA expression levels after exposure to conditioned media from LPS-stimulated adipocytes. *n≥7 mice per condition, pooled from 4 independent experiments. Statistical significance was calculated using (B,D-E) paired student’s t test or (F-G, I+J, K+L) by one-way ANOVA. *p < 0.05; **p < 0.01; ***p < 0.001; ****p < 0.0001. Illustration 3H was created with BioRender.com*.

Together, activated B cells in the presence of inflamed adipocytes demonstrate enhanced survival and differentiation. In turn, LPS-stimulated, activated B cells modulated the cytokine secretion profile of adipocytes, leading to increased secretion of MCP-1.

### B cell/adipocyte crosstalk promotes migration and polarisation of M2-like macrophages

The chemoattractant MCP-1 is involved in the recruitment of macrophages during inflammation. Given that LPS-stimulated adipocytes secrete MCP-1 into the culture supernatant (Fig. 2E, Suppl. Fig. 2E), and B cells even enhance this secretion (Fig. 3F), we next investigated the effect of adipocyte-released mediators on *in vitro* macrophage migration and polarisation. For this, we generated bone marrow-derived macrophages (BMDMs) *in vitro* (Fig. 3H). As macrophages themselves are highly sensitive to LPS stimulation, we used bone marrow isolated from C_3_H/HeJ mice, which do not respond to LPS (20-22). Consequently, macrophage polarization and activation was not influenced by LPS but by the soluble factors in the cell culture supernatants of LPS-stimulated adipocytes or B cell/adipocyte cocultures.

Using a transwell assay to study macrophage migration (Fig. 3H), we observed 2-fold higher numbers of migrated macrophages/mm^2^ in the presence of conditioned medium from LPS-activated adipocytes compared to unstimulated controls (Fig. 3I) after 4 hrs of culture. Adding anti-MCP-1 blocking antibodies to these cultures abrogated BMDM migration to baseline numbers (Fig. 3I). Comparing conditioned medium from LPS stimulated B cells, adipocytes or the B cell/adipocyte co-culture, we found that macrophage migration was further enhanced if cultured in conditioned medium of B cell/adipocyte co-cultures (Fig. 3J), correlating with the highest concentration of MCP-1 observed (Fig. 3F). Together, these results suggest that macrophage migration is mediated by MCP-1 secretion by LPS-simulated adipocytes, which is further enhanced in the presence of B cells.

As we have observed a high prevalence of M2-like macrophages in inflamed adipose tissue, we went on to test the influence of adipocyte conditioned media on macrophage polarisation (Fig. 3H). We found that M0 macrophages cultured for 24 hrs in the presence of conditioned medium from LPS-activated adipocytes demonstrated higher relative mRNA expression levels of iNOS (a marker for M1-like macrophage polarization, Suppl. Fig. 3D) as well as Arginase-1 (a marker for M2-like macrophage polarization, Suppl. Fig. 3E) compared to those incubated with conditioned medium from unstimulated adipocytes. Comparing conditioned medium from LPS stimulated B cells, adipocytes or the B cell/adipocyte co-culture, we found that relative mRNA expression levels of iNOS were comparable between all three conditioned media (Fig. 3K). Interestingly, mRNA expression levels of Arginase-1 were significantly elevated in macrophages exposed to condioned media of the LPS-stimulated B cell/adipocyte co-cultures, relative to macrophages cultured in condioned media of LPS-stimulated B cells or adipocytes alone (Fig. 3L). These results indicate an enhanced polarisation of M2-like macrophages in a environment primed by the B cell/adipocyte crosstalk.

### B cell/adipocyte crosstalk contributes to a lactate-rich, inflammatory microenvironment

In order to investigate which soluble mediators released by inflamed adipocytes could influence the polarization of M2-like, pro-fibrotic macrophages, we went on to analyse the culture supernatants of WAT-derived spheroids in the presence or absence of LPS using mass spectrometry. After LPS stimulation, we found an increase in orotidine and lactate concomitant with a reduction of citrate in the culture supernatant of adipocyte spheroids (Fig. 4A-D). Lactate, a key product of glycolysis, is produced when pyruvate is converted to lactate by the lactate dehydrogenase enzyme (LDHA. Enhanced levels of lactate concomitant with reduced levels of citrate were pointing at anaerobic glycolysis in hypoxic conditions. Analysing mRNA expression levels in adipocyte spheroids in the presence or absence of LPS indeed revealed a significant upregulation of the hypoxia marker hypoxia-inducible factor 1 alpha (HIF-1α) as well as NADPH oxidase 2 (Nox2), related to oxidative stress (Fig. 4E+F). Together, these findings indicate an hypoxic/inflamed state of LPS-stimulated adipocytes *in vitro*.

**Figure 4:**
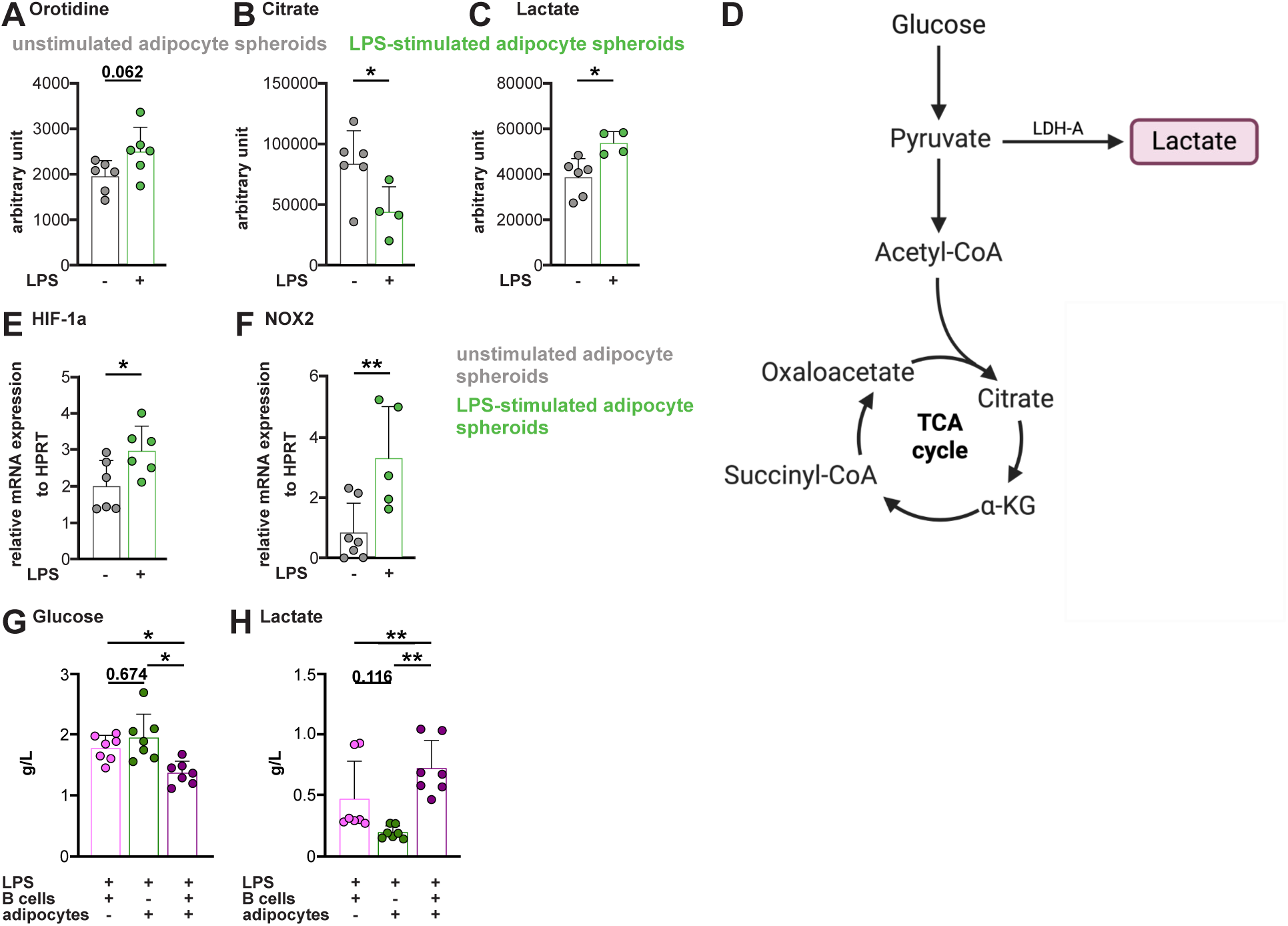
B cell/adipocyte crosstalk promotes a lactate-rich, inflammatory environment. (A-C) Non-targeted metabolomic analysis of culture supernatants from unstimulated and LPS-stimulated WAT spheroids. Relative quantification (arbitrary unit ± SD) of (A) orotidine, (B) citrate and (C) lactate levels. (D) Schematic overview of glucose metabolism and TCA cycle. Pyruvate is either converted to lactate via LDH-A or enters the TCA cycle through acetyl-CoA. (E+F) Relative mRNA expression (mean ± SD) of (E) HIF-1α and (F) NOX2 of non-stimulated and LPS stimulated WAT-derived spheroids, measured by qPCR. *Spheroids were generated from n≥4 WT mice, pooled from 3 independent experiments. Statistical significance was calculated using unpaired student’s t test.* (G+H) Glucose and lactate concentration were measured in supernatants of LPS-stimulated B cell, adipocytes or B cell/adipocyte co-cultures using the YSI analyser (g/L ± SD). *n≥7 mice per condition, pooled from 4 independent experiments. Statistical significance was calculated by one-way ANOVA. *p < 0.05; **p < 0.01;***p < 0.001; ****p < 0.0001. Illustration 4D was created with BioRender.com*.

Lactate has been shown to promote the differentiation of M2-like macrophages (23, 24). Having observed enhanced mRNA expression levels of Arginase-1 in our BMDM polarization assays in the presence of B cell/adipocyte conditioned medium (Fig. 3L), we next determined levels of lactate in the supernatants of the B cell/adipocyte co-cultures. Concomitant with a reduced levels of glucose (Fig. 4G), we found an increased concentration of lactate in the B cell/adipocyte conditioned medium compared to medium of B cells or adipocytes alone (Fig. 4H), which might enhance M2-like macrophage polarization.

### scRNAseq of mouse IBD-associated WAT and human CrF reveals high LDHA and HIF1A mRNA levels as well as a possible CCL2/CCR2-mediated macrophage recruitment

Results from our *in vitro* analysis suggested that the B cell/adipocyte crosstalk enhances the activation of both, B cells and adipocytes, leading to increased amounts of MCP-1 and lactate. Together, this lactate-rich, inflammatory environment promoted the migration and polarization of pro-fibrotic, M2-like macrophages *in vitro*. We next wanted to determine if we find evidence for the phenotype and the proposed mechanism also in *ex vivo* isolated mouse and human IBD-associated WAT tissue. First, we went on to analyse the phenotype of CD45+ leukocytes sorted from WAT of WT and *Wipf1^−/−^* mice using single-cell RNA sequencing (scRNAseq). Uniform manifold approximation and projection (UMAP) analyses showed that all major immune cell subsets were present in the WAT of WT and *Wipf1^−/−^* mice (Fig. 5A, Suppl. Fig. 4A-C). Using differential gene expression (DGE) analysis (hallmark gene set) on the B cell cluster revealed a significant enrichment of mRNA related to inflammation and pro-inflammatory cytokine response, especially IL-2/STAT5, TNF-α, and IFN-γ signalling in B cells isolated from inflamed WAT of *Wipf1^−/−^* mice compared to WAT of WT mice (Fig. 5B). Interestingly, differential pathway activity inference on the B cell cluster revealed a prominent hypoxia signature, in addition to many pathways related to cytokine signalling (including the JAK/STAT, Nfκ-B, PI3K and TGF-β pathways) in B cells isolated from WAT of *Wipf1^−/−^* mice (Suppl. Fig. 4D). In this line, ex vivo isolated B cells from inflamed WAT of *Wipf1^−/−^* mice demonstrated enhanced mRNA levels of *Ldha*, the enzyme responsible for converting pyruvate into lactate (Fig. 5C), as well as *Hif1a* (Fig. 5D). Together, these results suggested an enhanced lactate secretion in a hypoxic, activated state of B cells in IBD-associated WAT, supporting our *in vitro* findings.

**Figure 5:**
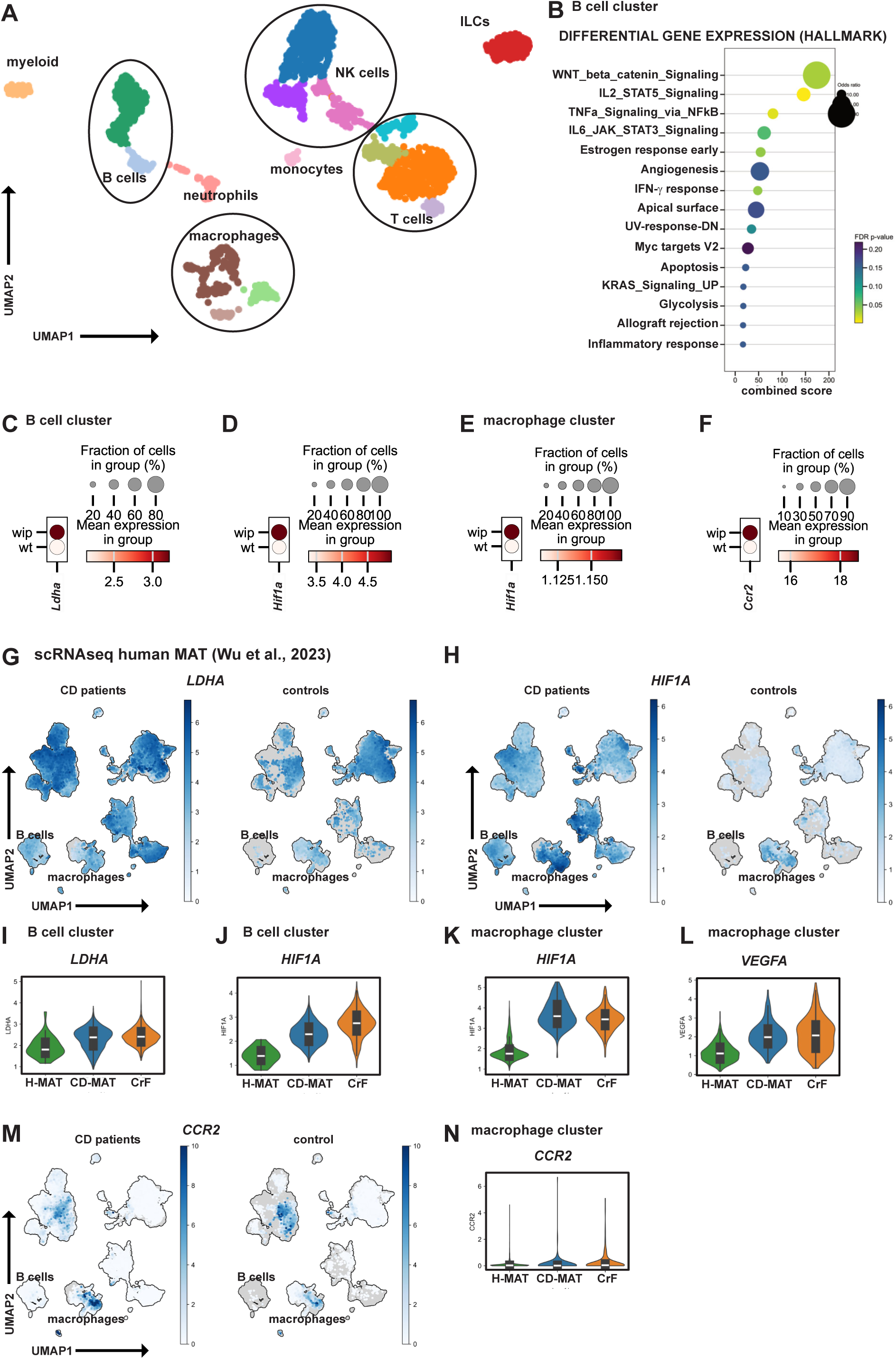
scRNAseq of mouse IBD-associated WAT reveals high *LDHA* and *HIF1A* mRNA levels as well as a possible CCL2/CCR2-mediated macrophage recruitment. (A) UMAP-representation of immune-cell clusters identified using scRNAseq analysis of CD45⁺ leukocytes isolated from WAT of WT and *Wipf1^−/−^* mice. (B) Overrepresentation analysis of functional terms (Hallmark gene sets) in B cell clusters. Dot size reflects odds ratio, color shows FDR-adjusted p-value, x-axis displays combined overrepresentation score. (C+D) mRNA expression levels of indicated genes in the B cell clusters split by genotype. Dot size indicates the proportion of cells expressing each gene within the group, while color intensity reflects the mean expression level. (E+F) Dot plot illustrating mRNA expression levels of indicated genes in the macrophage clusters split by genotype. Dot size indicates the proportion of cells expressing each gene within the group, while color intensity reflects the mean expression level. *(A-F) n≥2 mice per genotype.* (G+H, M) UMAP-representation of expression levels of (G) *LDHA*, (H) *HIF1A*, (M) *CCR2* in CrF/MAT of CD patients compared to control MAT. (I-L, N) Violin plots showing the distribution of (I) *LDHA*, (J+K) *HIF1A*, (L) *VEGFA,* (N) *CCR2* between CrF and uninvolved MAT (CD-MAT) from the same patient and MAT from healthy donor controls (H-MAT) in the indicated clusters (data extracted from Wu et al., 2023).

Using DGE analysis (hallmark gene set) on the macrophage cluster similarly revealed significant enrichment of mRNA related to inflammation and pro-inflammatory cytokine response, as well as hypoxia in macrophages isolated from inflamed WAT of *Wipf1^−/−^* mice (Suppl. Fig. 4E). In this line, we found prominent expression levels of *Hif1a* in macrophages isolated from adipose tissue of *Wipf1^−/−^* mice (Fig. 5E). Concomitantly, transcription factor activity inference demonstrated a higher *Hif1a* signature in *Wipf1^−/−^* macrophages (Suppl. Fig. 4F). Similarly, pathway activity inference demonstrated a higher hypoxia signature (Suppl. Fig. 4G), as well as pathways downstream of HIF-1α related to M2-like macrophage polarization, such as *Vegfa* (Suppl. Fig. 4G), all pointing at a macrophage response in a hypoxic/inflamed environment. Further correlating with our in vitro findings, we found that macrophages isolated from WAT of *Wipf1^−/−^* mice demonstrated higher expression levels of *Arginase1* (Suppl. Fig. 4H), which, together with the observed CD206 expression (Fig. 1H+I) points at an M2-like macrophage phenotype. Of note, macrophages isolated from WAT of *Wipf1^−/−^*mice demonstrated higher mRNA expression levels of C-C chemokine receptor 2 (*Ccr2*), indicating the capacity to migrate towards MCP-1 (Fig. 5 F). Collectively, our results provide evidence for an accumulation of M2-like macrophages with an inflammatory signature in hypoxic conditions, likely able to migrate toward an MCP-1 gradient in inflamed adipose tissue of *Wipf1^−/−^* mice.

We next re-analysed a scRNAseq dataset comparing CrF with uninvolved MAT (CD-MAT) from the same patient and MAT from healthy donor controls (H-MAT) (Fig. 5G-N, Suppl. Fig. 5A-C) (6). UMAP analysis revealed generally increased mRNA expression levels of *HIF1A* as well as *LDHA* (Fig. 5g+H) in MAT of CD patients. Similar to our findings analysing IBD-associated mouse WAT, B cells isolated from CrF of CD patients in this dataset showed enhanced mRNA expression levels of *LDHA* and *HIF1A* when compared to control MAT (Fig. 5I+J). In addition, macrophages from these datasets demonstrated enhanced levels of *HIF1A and VEGFA* (Fig. 5K+L). In addition, CCR2 mRNA levels were mainly upregulated in the monocyte/macrophage cluster in MAT of CD patients (Fig. 5M+N), similar to what we have observed comparing mouse inflamed WAT to control. In this human dataset, mesenchymal stroma cells (as well as other non-immune cells) displayed enhanced mRNA levels of MCP-1 (*CCL2*) (Suppl. Fig. 5C), as characterized in more detail in the original publication (6), supporting our in vitro findings of MCP-1 secretion by activated MAT-derived adipocyte spheroids (Fig. 2C, Suppl. Fig. 2B).

Collectively, our findings from scRNAseq analysis from mouse IBD-associated WAT and human CrF support the hypothesis that the B cell/adipocyte crosstalk fuels a lactate-rich, hypoxic, inflammatory environment, with enhanced production of MCP-1, which likely enables the recruitment and polarization of macrophages, thereby potentially driving IBD-associated WAT inflammation.

## Discussion

We here investigated how B cells shape the inflammatory responses in IBD-associated adipose tissue. We used high-parametric flow cytometry and showed an accumulation of B cells and plasma cells in human and mouse IBD-associated adipose tissue. Depleting B cells through anti-CD20 antibody treatment diminished M2-like macrophage accumulation in WAT in a mouse model of IBD. Using in vitro co-cultures, we found that the B cell/adipocyte crosstalk not only resulted in enhanced B cell activation but also led to elevated cytokine secretion by primary adipocytes, especially of MCP-1. This was in parts due to TNF-α secretion by activated B cells. Of note, supernatants of the B cell/adipocyte co-culture promoted M2-like macrophage polarisation in vitro, which correlated with enhanced levels of lactate. Concomitantly, scRNAseq analysis revealed that ex vivo isolated B cells from IBD-associated WAT of *Wipf1^−/−^*mice as well as from CrF of CD patients demonstrated enhanced mRNA levels of *Ldha*, indicating lactate production also in vivo. Interestingly, LPS-stimulated adipocyte spheroids in vitro, as well as macrophages and B cells isolated from human or mouse ex vivo displayed a hypoxic signature, pointing at a hypoxic environment in inflamed WAT. We thus hypothesize that the presence of B cells in IBD-associated WAT enhances MCP-1 secretion by adipocytes as well as the concentration of lactate in inflamed tissue. Elevated levels of MCP-1 will recruit more monocytes into the tissue, which will polarize into M2-like macrophages in a lactate-rich, hypoxic environment. Taken together, our results provide evidence for a crucial role of the B cell/adipocyte crosstalk to IBD-associated WAT inflammation.

While WAT inflammation related to obesity and aging is well studied, CD-associated CrF remains a poorly understood phenomenon. Hyperplasia of CrF in CD patients or inflamed WAT of experimental models is characterized by smaller, but increased numbers of adipocytes, as well as changes in their lipid metabolism (4, 6, 7, 25). Using Mid-infrared Optoacoustic Microscopy (MiROM), we recently found differences in the spectral distribution of adipocytes in IBD-related inflamed WAT of *Wipf1^−/−^*mice compared to WT tissue (5), also hinting at a difference in the lipid composition. Concomitantly, inflamed IBD-associated WAT, which contained both, adipocytes as well as immune cells, has been shown to express higher mRNA levels of inflammatory markers as well as chemoattractant molecules (26, 27). In order to better characterize adipocyte inflammation without the contribution of immune cells, we here investigated two different in vitro 3D culture methods of human or mouse adipocytes. We examined primary isolated mouse mature adipocytes in transwell cultures and developed human and mouse adipose tissue-derived 3D spheroid cultures. In contrast to the often used 3T3 pre-adipocyte model, those *in vitro* cultures have been shown to exhibit and maintain characteristic features (e.g. lipid metabolism) of *ex vivo* isolated WAT *in vitro* (28, 29). Of note, we found that adipocyte tissue-derived spheroids from CrF of CD patients readily secreted enhanced levels of pro-inflammatory cytokines without additional LPS stimulation, despite the prolonged culture condition. Furthermore, LPS stimulation of human and mouse adipose tissue-derived spheroids mimicked the cytokine secretion profile of unstimulated CrF-derived spheroids in our *in vitro* cultures. This is to the best of our knowledge the first characterization of a model of the pro-inflammatory potential of CrF adipocytes *in vitro*.

The secretion of pro-inflammatory cytokines of CrF-derived adipocyte spheroids suggested epigenetic imprinting of pre-adipocytes isolated from CrF of CD patients. Interestingly, during obesity, epigenetic regulation of adipogenesis as well as epigenetic transgenerational inheritance has been described (30). In CD, recent evidence pointed at metabolically primed fibroblasts isolated from stricturing CD compared to non-stricturing CD, correlating with distinct histone deacetylase expression profiles (31). Thus, pre-adipocytes isolated from the SVF of CrF of CD patients might be epigenetically primed to secrete pro-inflammatory cytokines, which might play a role in the recurrence of intestinal inflammation after bowel resection in CD patients. However, few studies investigated the influence of mesenteric removal on postoperative recurrence in CD patients (32, 33).

Although prominently present in CrF of CD patients, the role of B cells in CD-associated inflamed WAT has not been addressed so far. In contrast, during obesity or ageing, B cells have been demonstrated to be amongst the first immune cells to accumulate in inflamed WAT (34), and IgG antibodies have been demonstrated to be sufficient to cause insulin resistance in diet-induced obesity. Consequently, the absence of B cells and also IgG antibodies, either by B cell depletion therapy or using B cell deficient mice, prevented adipose tissue inflammation, WAT fibrosis and insulin resistance in diet-induced obesity or aging (13, 14, 35, 36). Our findings additionally highlight a central role of B cells in IBD-associated, non-obese, non-aged WAT inflammation. Depleting B cells through anti-CD20 antibody treatment at the onset of disease diminished M2-like macrophage and T cell accumulation in inflamed, IBD-associated mouse WAT. Mechanistically, our in vitro results suggest that activated B cells raise the concentration of TNF-a thereby promoting the secretion of MCP-1 by primary, mature, LPS-stimulated adipocytes. MCP-1 will subsequently lead to the recruitment of macrophages into inflamed adipose tissue. We cannot exclude that enhanced levels of IgG antibodies secreted by activated B cells might contribute to adipocyte activation and macrophage polarisation. Future experiments will address the role of IgG in IBD-associated WAT inflammation.

Our B cell/adipocyte co-culture assays also demonstrated enhanced expression of CD86 on activated B cells in the presence of adipocytes. We recently identified CD86 expression on activated B cells as a crucial factor for exacerbating pro-inflammatory cytokine production of intestinal CD4 T cells in *Wipf1^−/−^* mice (17). Blocking co-stimulatory signals mediated by CD86 through CTLA-4-Ig at the onset of disease diminished intestinal inflammation in our mouse model. It is thus tempting to speculate that CD86 expression on activated B cells might additionally contribute to an exacerbated inflammation through CD4 T cell co-stimulation in IBD-associated adipose tissue. Further experiments are needed to investigate the B cell/T cell interaction in mouse and human IBD-associated WAT.

Inflamed, IBD-associated adipose tissue has been linked to a hypoxic response, characterized by enhanced expression levels of *HIF1A* and lactate (7). In hypoxic conditions, cells increasingly switch to anaerobic glycolysis and convert pyruvate into lactate rather than entering the citric acid cycle after its conversion into acetyl-CoA. In addition, activated adipocytes also internally increase their oxygen consumption, leading to relative adipocyte hypoxia and lactate production (37, 38). Using targeted metabolomics of culture supernatants of LPS-stimulated adipose tissue-derived spheroids, we detected increased levels of a hypoxic marker, orotidine, as well extracellular lactate and diminished citrate levels. Furthermore, we found enhanced mRNA expression levels of *Hif1a* in LPS-stimulated, adipose tissue-derived spheroids, pointing at a hypoxic response in ambient oxygen conditions in a cell culture incubator. Thus, our 3D cell cultures suggested that adipose tissue hypoxia in IBD-associated, inflamed WAT might be initiated by adipocytes reacting to the presence of bacteria after barrier disruption. In addition, B cells might also contribute to enhanced local lactate concentrations in human and mouse inflamed IBD-associated adipose tissue. The effects of lactate on immune cells seem to be highly cell- and context-dependent (39). Lactate has been shown to promote the differentiation of anti-inflammatory, pro-angiogenic M2-like macrophages (23, 24).

During ageing, lactate seems to support B cell differentiation into age-associated B cells, characterized by autoantibody production, and the secretion of a senescence-associated secretory profile, such as IL-6 and TNF-α (40). Thus, lactate might also influence B cell differentiation in IBD-related inflamed adipose tissue. We cannot exclude that lactate secreted by LPS-stimulated adipocytes influences B cell survival and IgG plasmablast differentiation in our *in vitro* assays. Experiments are underway to address the exact mechanisms of how adipocytes promote B cell survival and influence B cell activation.

The chemoattractant MCP-1 is a key player in attracting monocytes/macrophages to inflamed obese or aged adipose tissue or IBD-associated WAT inflammation (27, 41-43). In addition, serum MCP-1 levels are associated with chronic inflammatory conditions during aging, obesity and autoimmune diseases. Of note, in UC patients, reduced serum levels of MCP-1 have been associated to successful therapy responses using anti-TNF-α antibodies (44). Recent data suggested a high and specific expression of the MCP-1 gene in a cluster with other fibrosis-associated genes as a genome-wide association signal for CD (6). Evidence from our re-analysis of this scRNAseq datasets from CD patients together with results from our *in vitro* and *in vivo* studies extend these findings by suggesting that an initial infiltration of B cells into adipose tissue might exacerbate the secretion of MCP-1 by adipocytes and contribute to a lactate-rich environment. As a result, monocytes/macrophages are recruited to IBD-related WAT. In addition to the presence of specific bacteria, the inflammatory, hypoxic and lactate-rich IBD-related adipose tissue microenvironment likely supports pro-fibrotic M2-like macrophage polarization. We recently described that depleting B cells through anti-CD20 antibody treatment at the onset of disease diminished intestinal inflammation in a mouse model of IBD (17). Our findings described here indicate that early B cell depletion might prevent or delay M2-like macrophage accumulation and thus the propagation of the vicious circle of IBD-related adipose tissue inflammation. Overall, our data suggests that the modulation of B cell activation during intestinal and adipose tissue inflammation might add to the therapeutic options for the treatment of IBD. Following the recent successes of B cell depletion therapy using CD19 CAR T cells in treating autoimmune diseases (45-47), we here present additional evidence supporting the key role of B cells in promoting inflammation across various types of tissues.

## Methods

### Sex as a biological variable

Human studies included both male and female participants as documented in Table 1, Supplemental Data - Table 1+2. Our study examined male and female animals, and similar findings are reported for both sexes.

### Mice

C57BL/6 WT mice were purchased from Charles River. Wipf1−/− mice were provided by Dr. Raif Geha (Boston’s Children Hospital, Boston, USA) (48) and backcrossed on C576BL/6 mice for at least 8 generations. C3H/HeJ mice were purchased from the Jackson Laboratory. All mice were housed and bred at the Specific Opportunist Pathogen Free (SOPF) animal facility at the TranslaTUM. Mice aged 6-16 weeks were used for all experiments. B cells were depleted from *Wipf1^−/−^* mice by i.p. administration of 250µg/ml anti-CD20 (SA271G2, Biolegend) or rat IgG2b, k Isotype control (RTK4530, Biolegend) at 6 weeks and 8 weeks of age, end analysis was performed at 10 weeks of age.

### Murine primary cell isolation and culture

Gonadal and mesenteric WAT were harvested from mice, minced, and enzymatically digested in 3 mL digestion medium (HBSS with Ca²⁺/Mg²⁺, 10 mM HEPES, 5% FCS (v/v), 1% penicillin-streptomycin (v/v), 3% BSA (w/v), 1 mg/ml collagenase D, 0.5 mg/ml dispase II, 0.25 mg/ml DNase I) for 40 min at 37°C with interval shaking. The suspension was filtered through 70 μm and 40 μm strainers (Greiner), separating tissue fragments and mature adipocytes from the stromal vascular fraction (SVF). Filters were rinsed with 5 mL WAT medium (advanced DMEM/F12, 1% GlutaMAX (v/v), 5% FCS (v/v), 1% penicillin-streptomycin (v/v)), and the SVF was pelleted by centrifugation at 1200 rpm for 10 min. The pellet was resuspended in WAT medium and plated in 10 cm dishes to expand adherent pre-adipocytes, with medium changes every 2–3 days until confluence. For mature adipocyte enrichment, tissue was minced, digested, filtered through a 100 μm strainer, and centrifuged at 50 g for 5 min at 4°C. Floating mature adipocytes were collected with wide-bore pipette tips, washed with WAT medium, and stored at 4°C. Lymphocytes were isolated from the SVF by Percoll gradient centrifugation (2200 rpm, 20 min, 4°C, 1 acceleration, 0 deceleration), collected at the 40/80% interface, and washed in B cell medium (RPMI-1640 + GlutaMax, 10mM HEPES, 10%FCS, 1x P/S and 0.05mM 2-Mercaptoethanol).

Splenic naïve B cells were purified using negative B cell isolation kits yielding enriched populations of 95%–98% (Miltenyi Biotec).

For the generation of bone marrow-derived macrophages (BMDMs), bone marrow isolated from the hind legs of C3H mice. After removing muscles and connective tissue, bone marrow was flushed from the the femur and tibia with PBS through a 70 μm filter. Cells were centrifuged (1200 rpm, 5 min, 4°C), and cultured in B cell medium with 40 ng/ml murine macrophage colony-stimulating factor (M-CSF) for 7 days to differentiate into M0 macrophages.

All cells were cultured at 37°C and 5% CO₂. Pre-adipocytes and mature adipocytes were maintained in WAT medium, while BMDM and B cell experiments used B cell medium. Adherent cells were split with 0.05% EDTA in PBS after reaching confluence. Viable cells were counted using trypan blue and a hemocytometer. For cryopreservation, cells were stored in FCS with 10% DMSO in liquid nitrogen.

### Isolation of human primary cells

MAT and CrF samples were collected by trained surgeons, washed with Ringer’s solution, weighed, and minced. Tissues were enzymatically digested for 1 hour at 37°C in HBSS buffer containing Ca²⁺/Mg²⁺, 10 mM HEPES, 5% FCS, 1% penicillin-streptomycin, 3% BSA, 1 mg/ml collagenase D, 0.5 mg/ml dispase II, and 0.25 mg/ml DNase I. The resulting suspension was filtered through a 100 μm strainer, and lymphocytes were purified through a Percoll (GE Healthcare) gradient (2200 rpm, 20 min, 4°C, 1 acceleration, 0 deceleration) harvested at the 40%/80% interface and washed in B cell medium (RPMI-1650 + GlutaMax, 10 mM HEPES, 10% FCS, 1% P/S, and 0.05 mM 2-mercaptoethanol).

### Flow cytometric analysis

Human and mouse single cell suspensions were stained for viability using the Zombie Aqua fixable viability kit (1:1000 BioLegend) and fixed with 4% paraformaldehyde (PFA, VWR international). Cells were blocked with anti-mouse CD16/32 (93, 1:200, BioLegend) and CD16.2 (9E9, 1:200, BioLegend) followed by staining with a combination of antibodies (see Supplemental Data - Table 3) for 20 min on ice, washed, and analysed on a LSR Fortessa cytometer (BD Biosciences) using FlowJo software.

### Human and mouse adipose-tissue-derived spheroid cultures

Pr-eadipocytes isolated from human (3×10⁴ cells/well) or murine (5×10⁴ cells/well) adipose tissue were seeded into cell-repellent 96-well U-bottom plates (Greiner) to form spheroids over 1–2 days. Spheroids were differentiated into adipocytes using WAT medium with 5 μM dexamethasone, 0.5 μg/mL insulin, 0.5 mM IBMX, and 1 μM rosiglitazone for 2 days, then maintained in WAT medium with 0.5 μg/mL insulin for ∼3 weeks. Three mature spheroids were pooled and stimulated with 5 μg/mL LPS for 2 days for mRNA and cytokine analysis.

### B cell/adipocyte co-cultures

A transwell culture system was used for co-culture experiments. Isolated mature adipocytes (20 μl) were pipetted onto inverted Transwells (0.4 μm, Sarstedt) and placed in 24-well plates with 0.5–1 ml B cell medium. After 4 hours, 8×10⁵ purified *Wipf1^−/−^* B cells were added to each membrane. Co-cultures were stimulated with 5 μg/ml LPS for 4 days.

For neutralization assays, 5 μg/ml neutralizing antibody (anti-IL6, anti-IL10, anti-TNFα, or anti-IFNγ) was added at day 0 and day 2. B cells were collected for FACS analysis, and supernatants were used for cytokine profiling (LEGENDplex multiplex assays), monocyte migration and differentiation assays, and lactate concentration measurements (YSI Model 2950 Analyzer).

### Macrophage assays

Macrophage migration was assessed using 8 μm Transwell inserts (Sarstedt). Supernatants from Transwell co-culture experiments were used as chemoattractants in 24-well plates. 1×10⁵ M0 macrophages were added to the upper chamber and incubated for 4 hours at 37°C. Non-migrated cells were removed, and migrated cells on the underside were stained with Hoechst for 10 minutes and counted at three random sites using a fluorescence microscope (EVOS FL). Cell counts were normalized to the insert surface area (mm²).

For macrophage activation/polarization with Transwell co-culture supernatants, 1×10^5^ M0 macrophages were cultured with an equal 100 μl co-culture medium volume for 12-18 hours at 37°C. Cells were lysed in RLT buffer for qPCR analysis.

### RNA extraction, reverse transcription, quantitative polymerase chain reaction (qPCR)

Total RNA was isolated from 1×10⁵ polarized macrophages or pooled adipose spheroids using the RNeasy Micro Kit (Qiagen), with tissue disrupted in RLT buffer. 10 μl RNA was reverse transcribed to cDNA (High-Capacity cDNA Reverse Transcription Kit, Applied Biosystems). qPCR was performed with Takyon No ROX SYBR 2x MasterMix (Eurogentec) and specific primers (Supplemental Data - Table 4) on a Light Cycler 480II (Roche). Relative gene expression was calculated using the 2^−ΔCt^ method with *Hprt* (mouse) as housekeeping genes.

### Bacterial DNA extraction and quantification

Bacterial genomic DNA was extracted from 100 mg mWAT using the DNeasy PowerSoil Kit (Qiagen), with tissue disruption in PowerBead tubes. Samples were processed per the manufacturer’s protocol, and 100 μl of eluted gDNA was stored at –20°C. DNA concentration and quality were assessed with a NanoDrop spectrophotometer (Thermo Scientific). Quantitative PCR (qPCR) was used to measure 16S rDNA copy number in adipose tissue (Jimeno et al., 2018). A standard curve was generated from serially diluted plasmid DNA containing the V3-V4 region of 16S rDNA. Bacterial concentration was calculated from Ct values and expressed as 16S rDNA copies/μl DNA extract. Each qPCR reaction included 2 μl of standard or eluted gDNA.

### Multiplex assay

Cytokines and immunoglobulins in culture supernatants were quantified using LEGENDplex multiplex assays (BioLegend) per the manufacturer’s instructions. For murine samples, we used the Mouse Inflammation (13-plex) or Immunoglobulin Isotyping (6-plex) panels; for human samples we used the Human Adipokine Panel (13-plex).

### Non-targeted metabolomics

Non-targeted metabolomics was performed on extracts from spheroid culture supernatants. Polar metabolites were extracted with ice-cold 80% methanol, vacuum-concentrated, and stored at –70°C until measurement. For LC-MS analyzes an Agilent 1290 Infinity II UHPLC inline with a Bruker Impact II QTOF-MS operating in negative ion mode was used. Scan range was from 20 to 1050 Da. Mass calibration was performed at the beginning of each run. LC separation was on a Waters Atlantis Premier BEH ZHILIC column (100 x 2.1 mm, 1.7 µm particles), buffer A was 20 mM ammonium carbonate and 5 µM medronic acid in milliQ H_2_O and buffer B was 90:10 acetonitrile:buffer A and the solvent gradient was from 95% to 55% buffer B over 14 minutes. Flow rate was 180 µL/min, column temperature was 35 °C, autosampler temperature was 5 °C and injection volume was 3 µL. Data processing including detection, deconvolution, and identification of features was carried out using Metaboscape 2023 (Bruker). At least three negative controls were run per experiment.

### Whole-mount Immunofluorescence

For immunofluorescence, WAT was processed as previously described (18). Tissue was fixed (IC fixation buffer, Thermo Fisher FB001; 1 h, RT), blocked overnight (1% (v/v) FCS, 1% (v/v) mouse serum, and 0.3% (v/v) Triton X-100 in PBS) at 37°C, and stained in blocking buffer with BV421 anti-CD163, AF488 anti-MHC-II, AF594 anti-CD11b, and AF647 anti-CD206 (all 1:200) for 72 h at 37°C. After washing (0.2 % (v/v) Triton X-100 and 0.5 % (v/v) 1-thioglycerol in PBS; 24 h, 37°C), tissue was dehydrated in graded isopropanol (30% - 50% - 70% - 100%, Sigma-Aldrich) and cleared with ethyl cinnamate (Sigma-Aldrich #W243000). Imaging was performed using an inverted Leica TCS SP8 confocal microscope equipped with a white light laser and HyD photodetectors, utilizing an HC PL APO CS2 20×/0.75 IMM objective at a zoom factor of 1 (Leica Microsystems). Image deconvolution was carried out with the LIGHTNING module of the Leica Application Suite X (Leica Microsystems). Subsequent image analysis and contrast adjustments were completed in Imaris (Bitplane, version 9.5.1). Immune cells were segmented by generating binary masks with the Surface Creation Wizard in Imaris, and quantitative data for these objects were exported for cell quantification.

### Single-cell RNA sequencing

For single cell sequencing, CD45+ immune cells from WAT were sorted on the BD Aria. Single cell suspension and library preparation was performed with 20.000 of these cells on the 10x genomics platform and following the Chromium Next GEM Single Cell 3’ Reagent Kit User Guide Revision E. Illumina Next Generation Sequencing was performed using an Illumina NovaSeq 6000 sequencing system by the Helmholtz Munich Core Facility Genomics, including raw data quality control, alignment and mapping using Cell Ranger. The resulting data was then quality controlled and analysed using the scanpy toolkit (49) and decoupleR (50) (Supplemental File).

### Human MAT single-cell RNA sequencing re-analysis

Annotated gene expression matrix of immune cells (6), were loaded, processed and re-analyzed using the scanpy toolkit (ver. 1.11.2, (51), SCR_01813999), scvi-tools (ver. 1.3.1, SCR_026673), and other common Python (ver. 3.13, SCR_008394) toolboxes.

Low-quality observations, filtered individually per sample based on observed distribution by number of expressed unique genes (minimum: 500–1250, maximum: 3500–7500), total read counts (maximum: 11500-50000), and percentage of mitochondrial genes (maximum.: 15%– 18%) and genes expressed by fewer than three cells were removed. For sample integration highly variable genes (flavor: “seurat_v3”, n_top_genes: 15000, layer: “counts”, batch_key: “sample_ID”, subset: True) were calculated and a single cell Variational Inference (scVI (52)) model (layer: “counts”, batch_key: “sample_ID”, n_latent: 30, n_layers: 2) was trained (max_epochs: 400, early_stopping: True, early_stopping_patience: 20, early_stopping_monitor: “elbo_validation”). Next, the neighborhood graph (nearest neighbors: 50) based on the scVI model’s latent space representation, Uniform Manifold Approximation and Projection (UMAP; SCR_018217103; min. distance: 0.4, neg. edge sample rate: 12), and initial Leiden clustering (ver. 0.10.2; resolution: 1.0) were computed as indicated and used as basis for cell type annotations in combination with computed marker genes (method: “wilcoxon”, corr_method: “bonferroni”). For displays of gene expression, normalized values were extracted from the integration model (library_size: 1e4), log1p transformed, and scaled (max_value: 10, zero_center: False).

### Statistics

GraphPad Prism 10.1. was used to generate graphs and for all statistical analyses other than those performed on the RNA sequencing data. Figure legends detail the number of experimental replicates and n values. Statistical analysis used paired or unpaired two-sided Student’s t-tests or one-way ANOVA. Error bars represent mean ± SD unless otherwise indicated in the figure legends. *p < 0.05; **p < 0.01; ***p < 0.001; ****p < 0.0001.

### Study approval

Informed, written consent was obtained from all patients during their pre-operative visit, with prior approval of the local ethics committee of the Faculty of Medicine of the Technical University of Munich, including two-fold pseudonymization for patient tissue (TUM; #1926/2007, #5428/12 and 2022-297-S22-KH). Fresh mesenteric adipose tissue samples were collected adjacent to inflamed/fibrotic and non-involved bowel segments from CD patients undergoing intestinal resections. Tissue sampling was performed in accordance with the regulations of the tissue bank of the TUM and Klinikum rechts der Isar, Munich (MTBIO). All experiments involving mice were conducted in compliance with federal and institutional guidelines, and animal protocols were approved by the Government of Upper Bavaria (Regierung von Oberbayern).

## Data availability

All data supporting the findings of this study are available within the article or the supplemental materials and from the corresponding author upon reasonable request. Raw scRNA-seq data (FASTQ format) are available from the XX repositories (XX accessions:xx).

## Author contributions

Conceptualization: SJK

Methodology: MG, JMH, IG, AB, MW, JH

Investigation: MG, JMH, IG, AB

Visualization: SJK, MG

Funding acquisition: SJK, PAN, JR

Project administration: SJK

Supervision: SJK

Writing – original draft: SJK

Writing – review & editing: SJK

## Acknowledgments

We thank Raif Geha for providing *Wipf1^−/−^*mice. We further thank Martin Klingenspor and all members of his group, Lola de la Fuentes and Agnes Huber for valuable expertise and providing reagents. We thank the biological resource unit for animal husbandry (ZPF of the MRI at TUM) and the core facility for flow cytometry at the TranslaTUM (Linda Bachmann), the Helmholtz Munich Core Facility Genomics and Daniel Strobl, as well as the Metabolomics and FACS core facility at the MPI-IE for excellent support. We further thank all members of the Rheumatology and Immunology lab at the Medical University Graz for scientific discussions and support.

## References

1. Ha CWY, Martin A, Sepich-Poore GD, Shi B, Wang Y, Gouin K, et al. Translocation of Viable Gut Microbiota to Mesenteric Adipose Drives Formation of Creeping Fat in Humans. Cell. 2020;183(3):666–83 e17.

2. Peyrin-Biroulet L, Chamaillard M, Gonzalez F, Beclin E, Decourcelle C, Antunes L, et al. Mesenteric fat in Crohn’s disease: a pathogenetic hallmark or an innocent bystander? Gut. 2007;56(4):577–83.

3. Crohn BB, Ginzburg L, Oppenheimer GD. Landmark article Oct 15, 1932. Regional ileitis. A pathological and clinical entity. By Burril B. Crohn, Leon Ginzburg, and Gordon D. Oppenheimer. JAMA. 1984;251(1):73–9.

4. Liu W, Mao R, Nga Le TH, West G, Varadharajan V, Banerjee R, et al. Creeping Fat-Derived Free Fatty Acids Induce Hyperplasia of Intestinal Muscularis Propria Muscle Cells: A Novel Link Between Fat and Intestinal Stricture Formation in Crohn’s Disease. Gastroenterology. 2025;168(3):508–24.

5. Ko V, Goess MC, Scheel-Platz L, Yuan T, Chmyrov A, Justel D, et al. Fast histological assessment of adipose tissue inflammation by label-free mid-infrared optoacoustic microscopy. Npj Imaging. 2023;1(1):3.

6. Wu F, Wu F, Zhou Q, Liu X, Fei J, Zhang D, et al. A CCL2(+)DPP4(+) subset of mesenchymal stem cells expedites aberrant formation of creeping fat in humans. Nat Commun. 2023;14(1):5830.

7. Zuo L, Li Y, Zhu W, Shen B, Gong J, Guo Z, et al. Mesenteric Adipocyte Dysfunction in Crohn’s Disease is Associated with Hypoxia. Inflamm Bowel Dis. 2016;22(1):114–26.

8. Kredel LI, Batra A, Stroh T, Kuhl AA, Zeitz M, Erben U, et al. Adipokines from local fat cells shape the macrophage compartment of the creeping fat in Crohn’s disease. Gut. 2013;62(6):852–62.

9. Batra A, Pietsch J, Fedke I, Glauben R, Okur B, Stroh T, et al. Leptin-dependent toll-like receptor expression and responsiveness in preadipocytes and adipocytes. Am J Pathol. 2007;170(6):1931–41.

10. Paul G, Schaffler A, Neumeier M, Furst A, Bataillle F, Buechler C, et al. Profiling adipocytokine secretion from creeping fat in Crohn’s disease. Inflamm Bowel Dis. 2006;12(6):471–7.

11. Yamamoto K, Kiyohara T, Murayama Y, Kihara S, Okamoto Y, Funahashi T, et al. Production of adiponectin, an anti-inflammatory protein, in mesenteric adipose tissue in Crohn’s disease. Gut. 2005;54(6):789–96.

12. da Silva FAR, Pascoal LB, Dotti I, Setsuko Ayrizono ML, Aguilar D, Rodrigues BL, et al. Whole transcriptional analysis identifies markers of B, T and plasma cell signaling pathways in the mesenteric adipose tissue associated with Crohn’s disease. J Transl Med. 2020;18(1):44.

13. DeFuria J, Belkina AC, Jagannathan-Bogdan M, Snyder-Cappione J, Carr JD, Nersesova YR, et al. B cells promote inflammation in obesity and type 2 diabetes through regulation of T-cell function and an inflammatory cytokine profile. Proceedings of the National Academy of Sciences of the United States of America. 2013;110(13):5133–8.

14. Winer DA, Winer S, Shen L, Wadia PP, Yantha J, Paltser G, et al. B cells promote insulin resistance through modulation of T cells and production of pathogenic IgG antibodies. Nat Med. 2011;17(5):610–7.

15. Camell CD, Gunther P, Lee A, Goldberg EL, Spadaro O, Youm YH, et al. Aging Induces an Nlrp3 Inflammasome-Dependent Expansion of Adipose B Cells That Impairs Metabolic Homeostasis. Cell Metab. 2019;30(6):1024–39 e6.

16. Hagglof T, Vanz C, Kumagai A, Dudley E, Ortega V, Siller M, et al. T-bet(+) B cells accumulate in adipose tissue and exacerbate metabolic disorder during obesity. Cell Metab. 2022;34(8):1121–36 e6.

17. Gadjalova I, Heinze JM, Goess MC, Hofmann J, Buck A, Weber MC, et al. B cell-mediated CD4 T-cell costimulation via CD86 exacerbates pro-inflammatory cytokine production during autoimmune intestinal inflammation. Mucosal Immunol. 2024;17(1):67–80.

18. Hofmann J, Gadjalova I, Mishra R, Ruland J, Keppler SJ. Efficient Tissue Clearing and Multi-Organ Volumetric Imaging Enable Quantitative Visualization of Sparse Immune Cell Populations During Inflammation. Front Immunol. 2020;11:599495.

19. Maniyadath B, Zhang Q, Gupta RK, Mandrup S. Adipose tissue at single-cell resolution. Cell Metab. 2023;35(3):386–413.

20. Poltorak A, He X, Smirnova I, Liu MY, Van Huffel C, Du X, et al. Defective LPS signaling in C3H/HeJ and C57BL/10ScCr mice: mutations in Tlr4 gene. Science. 1998;282(5396):2085–8.

21. Qureshi ST, Lariviere L, Leveque G, Clermont S, Moore KJ, Gros P, et al. Endotoxin-tolerant mice have mutations in Toll-like receptor 4 (Tlr4). The Journal of experimental medicine. 1999;189(4):615–25.

22. Ding A, Hwang S, Lander HM, Xie QW. Macrophages derived from C3H/HeJ (Lpsd) mice respond to bacterial lipopolysaccharide by activating NF-kappa B. J Leukoc Biol. 1995;57(1):174–9.

23. Colegio OR, Chu NQ, Szabo AL, Chu T, Rhebergen AM, Jairam V, et al. Functional polarization of tumour-associated macrophages by tumour-derived lactic acid. Nature. 2014;513(7519):559–63.

24. Noe JT, Rendon BE, Geller AE, Conroy LR, Morrissey SM, Young LEA, et al. Lactate supports a metabolic-epigenetic link in macrophage polarization. Sci Adv. 2021;7(46):eabi8602.

25. Peyrin-Biroulet L, Gonzalez F, Dubuquoy L, Rousseaux C, Dubuquoy C, Decourcelle C, et al. Mesenteric fat as a source of C reactive protein and as a target for bacterial translocation in Crohn’s disease. Gut. 2012;61(1):78–85.

26. Mustain WC, Starr ME, Valentino JD, Cohen DA, Okamura D, Wang C, et al. Inflammatory cytokine gene expression in mesenteric adipose tissue during acute experimental colitis. PLoS One. 2013;8(12):e83693.

27. Olivier I, Theodorou V, Valet P, Castan-Laurell I, Ferrier L, Eutamene H. Modifications of mesenteric adipose tissue during moderate experimental colitis in mice. Life Sci. 2014;94(1):1–7.

28. Harms MJ, Li Q, Lee S, Zhang C, Kull B, Hallen S, et al. Mature Human White Adipocytes Cultured under Membranes Maintain Identity, Function, and Can Transdifferentiate into Brown-like Adipocytes. Cell Rep. 2019;27(1):213–25 e5.

29. Taylor J, Sellin J, Kuerschner L, Krahl L, Majlesain Y, Forster I, et al. Generation of immune cell containing adipose organoids for in vitro analysis of immune metabolism. Sci Rep. 2020;10(1):21104.

30. Pant R, Firmal P, Shah VK, Alam A, Chattopadhyay S. Epigenetic Regulation of Adipogenesis in Development of Metabolic Syndrome. Front Cell Dev Biol. 2020;8:619888.

31. Lewis A, Humphreys DT, Pan-Castillo B, Berti G, Felice C, Gordon H, et al. Epigenetic and Metabolic Reprogramming of Fibroblasts in Crohn’s Disease Strictures Reveals Histone Deacetylases as Therapeutic Targets. J Crohns Colitis. 2024;18(6):895–907.

32. Topala M, Martinekova P, Rancz A, Veres DS, Lenti K, Miheller P, et al. To cut or not to cut? Extended mesenteric excision during intestinal resection does not impact the postoperative recurrence nor the postoperative complications in Crohn’s disease: a systematic review and meta-analysis. Tech Coloproctol. 2025;29(1):79.

33. Coffey JC, O’Leary DP, Kiernan MG, Faul P. The mesentery in Crohn’s disease: friend or foe? Curr Opin Gastroenterol. 2016;32(4):267–73.

34. Duffaut C, Galitzky J, Lafontan M, Bouloumie A. Unexpected trafficking of immune cells within the adipose tissue during the onset of obesity. Biochem Biophys Res Commun. 2009;384(4):482–5.

35. Ying W, Wollam J, Ofrecio JM, Bandyopadhyay G, El Ouarrat D, Lee YS, et al. Adipose tissue B2 cells promote insulin resistance through leukotriene LTB4/LTB4R1 signaling. J Clin Invest. 2017;127(3):1019–30.

36. Yu L, Wan Q, Liu Q, Fan Y, Zhou Q, Skowronski AA, et al. IgG is an aging factor that drives adipose tissue fibrosis and metabolic decline. Cell Metab. 2024;36(4):793–807 e5.

37. Krycer JR, Quek LE, Francis D, Fazakerley DJ, Elkington SD, Diaz-Vegas A, et al. Lactate production is a prioritized feature of adipocyte metabolism. The Journal of biological chemistry. 2020;295(1):83–98.

38. Lee YS, Kim JW, Osborne O, Oh DY, Sasik R, Schenk S, et al. Increased adipocyte O2 consumption triggers HIF-1alpha, causing inflammation and insulin resistance in obesity. Cell. 2014;157(6):1339–52.

39. Llibre A, Kucuk S, Gope A, Certo M, Mauro C. Lactate: A key regulator of the immune response. Immunity. 2025;58(3):535–54.

40. Romero M, Miller K, Gelsomini A, Garcia D, Li K, Suresh D, et al. Immunometabolic effects of lactate on humoral immunity in healthy individuals of different ages. Nat Commun. 2024;15(1):7515.

41. Kanda H, Tateya S, Tamori Y, Kotani K, Hiasa K, Kitazawa R, et al. MCP-1 contributes to macrophage infiltration into adipose tissue, insulin resistance, and hepatic steatosis in obesity. J Clin Invest. 2006;116(6):1494–505.

42. Karagiannides I, Kokkotou E, Tansky M, Tchkonia T, Giorgadze N, O’Brien M, et al. Induction of colitis causes inflammatory responses in fat depots: evidence for substance P pathways in human mesenteric preadipocytes. Proceedings of the National Academy of Sciences of the United States of America. 2006;103(13):5207–12.

43. Yu R, Kim CS, Kwon BS, Kawada T. Mesenteric adipose tissue-derived monocyte chemoattractant protein-1 plays a crucial role in adipose tissue macrophage migration and activation in obese mice. Obesity (Silver Spring). 2006;14(8):1353–62.

44. Magnusson MK, Strid H, Isaksson S, Bajor A, Lasson A, Ung KA, et al. Response to infliximab therapy in ulcerative colitis is associated with decreased monocyte activation, reduced CCL2 expression and downregulation of Tenascin C. J Crohns Colitis. 2015;9(1):56–65.

45. Schett G, Nagy G, Kronke G, Mielenz D. B-cell depletion in autoimmune diseases. Ann Rheum Dis. 2024.

46. Muller F, Taubmann J, Bucci L, Wilhelm A, Bergmann C, Volkl S, et al. CD19 CAR T-Cell Therapy in Autoimmune Disease - A Case Series with Follow-up. N Engl J Med. 2024;390(8):687–700.

47. Mackensen A, Muller F, Mougiakakos D, Boltz S, Wilhelm A, Aigner M, et al. Anti-CD19 CAR T cell therapy for refractory systemic lupus erythematosus. Nat Med. 2022;28(10):2124–32.

48. Anton IM, de la Fuente MA, Sims TN, Freeman S, Ramesh N, Hartwig JH, et al. WIP deficiency reveals a differential role for WIP and the actin cytoskeleton in T and B cell activation. Immunity. 2002;16(2):193–204.

49. Heumos L, Schaar AC, Lance C, Litinetskaya A, Drost F, Zappia L, et al. Best practices for single-cell analysis across modalities. Nat Rev Genet. 2023;24(8):550–72.

50. Badia IMP, Velez Santiago J, Braunger J, Geiss C, Dimitrov D, Muller-Dott S, et al. decoupleR: ensemble of computational methods to infer biological activities from omics data. Bioinform Adv. 2022;2(1):vbac016.

51. Wolf FA, Angerer P, Theis FJ. SCANPY: large-scale single-cell gene expression data analysis. Genome Biol. 2018;19(1):15.

52. Lopez R, Regier J, Cole MB, Jordan MI, Yosef N. Deep generative modeling for single-cell transcriptomics. Nat Methods. 2018;15(12):1053–8.

