## Supplemental FIle for "B cell/adipocyte crosstalk drives pro-fibrotic macrophage migration and polarisation in Crohn’s disease-associated, inflamed adipose tissue"

### Supplemental Figures

Supplemental Figure 1 – related to Figure 1

Supplemental Figure 2 – related to Figure 2

Supplemental Figure 3 – related to Figure 3

Supplemental Figure 4 – related to Figure 5

Supplemental Figure 5 – related to Figure 5

### Supplemental Tables

Supplemental Table 1: patients` characteristics – flow

Supplemental Table 2: patients` characteristics - spheroids

Supplemental Table 3: list of antibodies

Supplemental Table 4: list of oligonucleotides

### Suppl. Fig. 1 related to Fig. 1

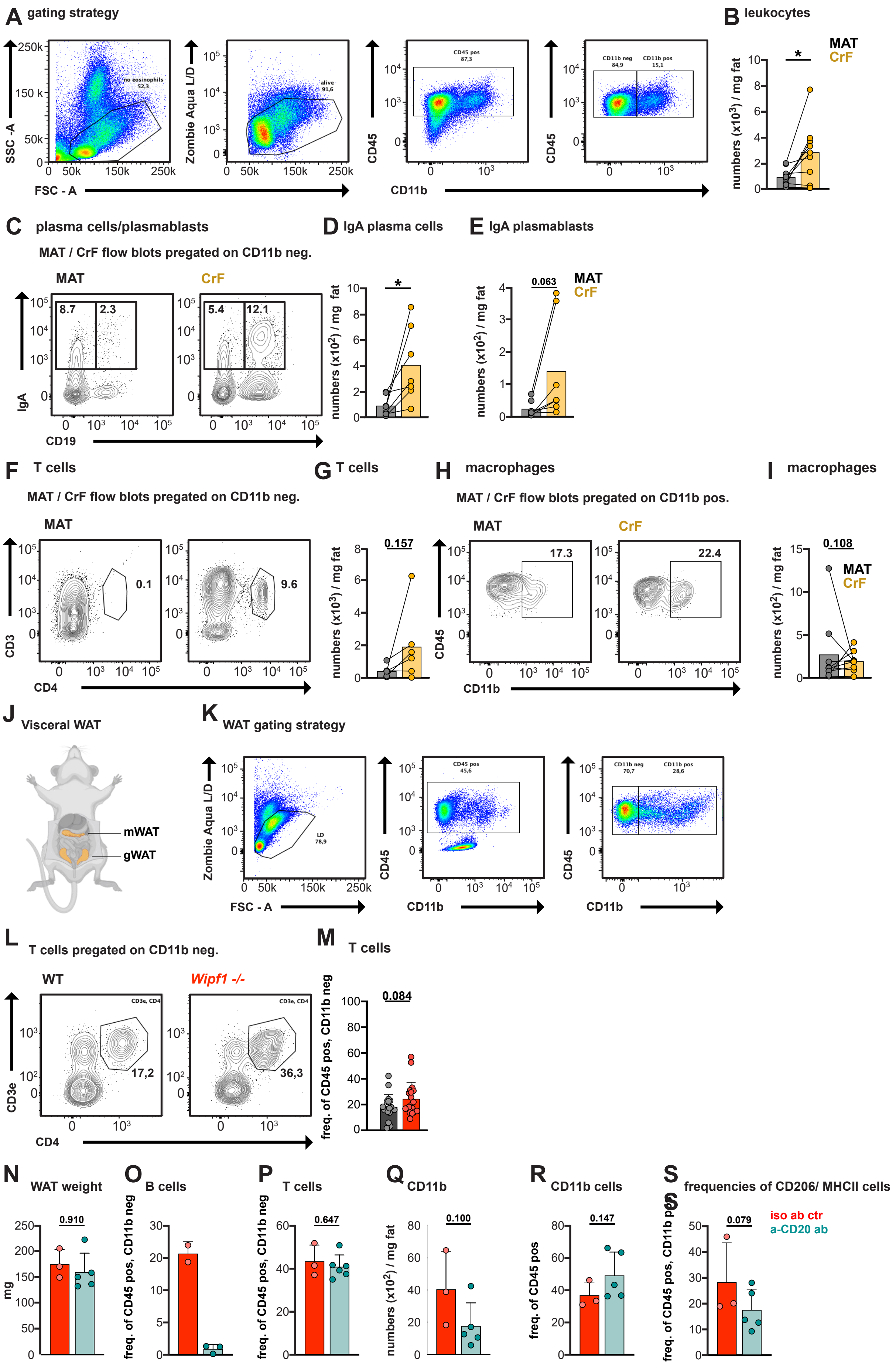

**Suppl. Fig. 1:**

(A) Gating strategy for leukocytes in single cell suspensions isolated from MAT and CrF of CD patients. (B) Total number of living leukocytes ($CD45^+$) per mg fat tissue from MAT and CrF. (C) Representative plots of IgA plasma cells and IgA plasmablasts, gated on $CD45^+$, $CD11b^-$. (D) Total cell number of IgA plasma cells ($CD19^-$, $IgA^+$) per mg fat and (E) IgA plasmablasts ($CD19^+$, $IgA^+$) per mg fat. (F) Representative plots of T cells, gated on $CD45^+$, $CD11b^-$. (G) Total cell number of T cells per mg fat. (H) Representative plots of macrophages, gated on $CD45^+$. (I) Total cell number of macrophages per mg fat. Paired samples from MAT and CrF were collected from each patient ($n \geq 7$); bars indicate the mean.

(J) Schematic depicting the regions of visceral white adipose tissue (mWAT and gWAT) in mice collected for analysis. (K) Flow cytometry gating strategy used to assess the frequencies of (L+M) CD3+, CD4+ T cells in single cell suspensions prepared from visceral WAT. Data represent $n \geq 10$ mice per genotype, pooled from multiple independent experiments.

(N) WAT weight was recorded following tissue harvest from experimental mice. Frequencies of (O) B cells, (P) T cells, (R) CD11b cells and (S) CD206/MHCII cells were measured in WAT following anti-CD20 antibody treatment. (Q) Total numbers of CD11b cells quantified in WAT after anti-CD20 antibody treatment. $n \geq 3$ mice per genotype, pooled from 2 independent experiments. Statistical significance was determined using a paired student´s *t* test. *$p < 0.05$; **$p < 0.01$; ***$p < 0.001$; ****$p < 0.0001$. Illustration 1J was created with BioRender.com.

### Suppl. Fig. 2 - related to Fig. 2

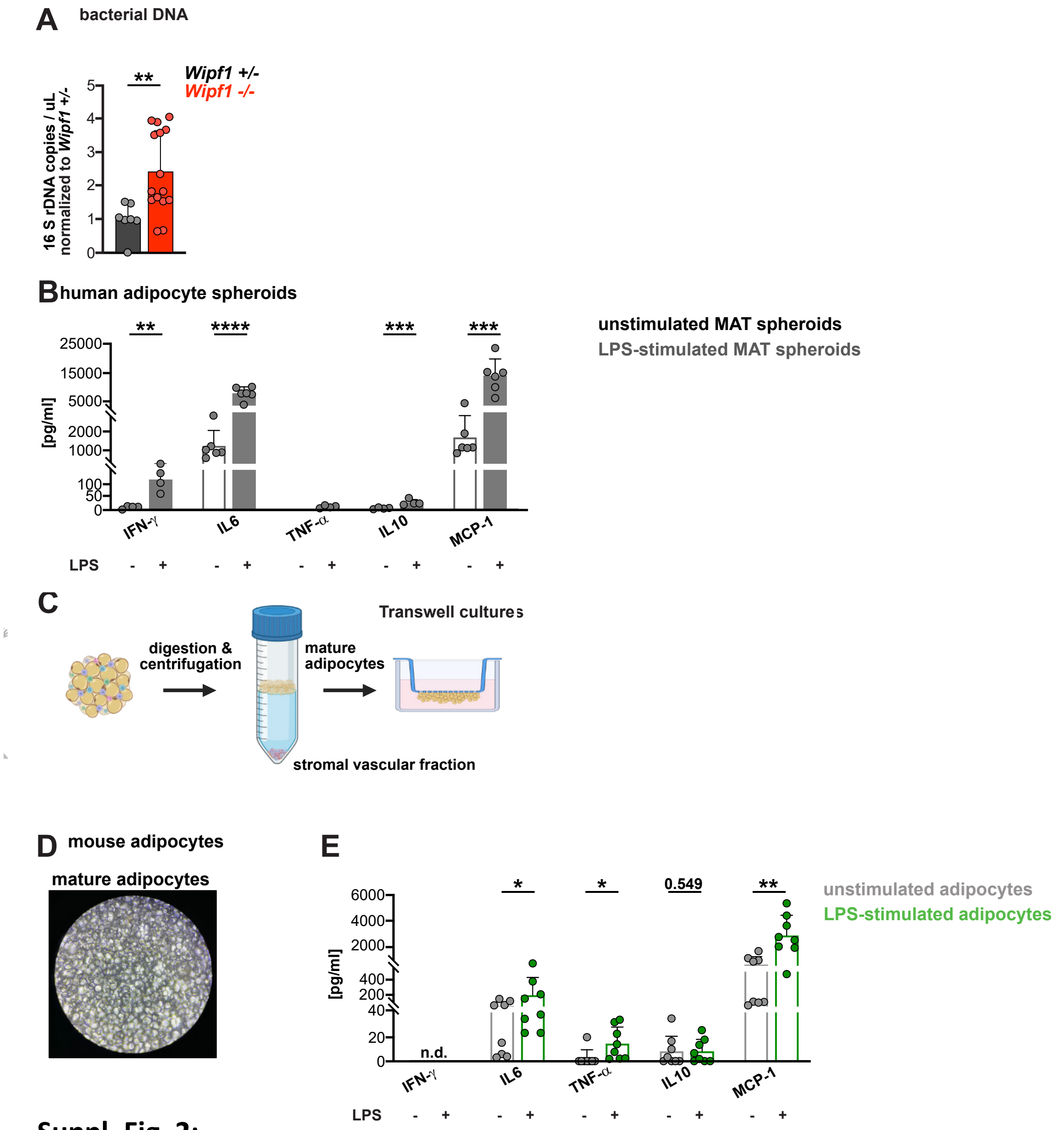


**Suppl. Fig. 2:**

(A) Bacterial DNA (16S rDNA copies/µL of DNA extract) in WAT of *Wipf1*$^{+/-}$ and *Wipf1*$^{-/-}$ mice. n≥4 mice per genotype. Data is presented as fold change ± SD relative to control *Wipf1*$^{+/-}$ mice. Statistical significance was calculated using unpaired student´s *t* test. (B) Cytokine secretion pattern of differentiated MAT-derived adipocyte spheroids cultured under non-stimulated and LPS stimulated conditions was measured using a multiplex immunoassay (Legenplex) on collected culture supernatants. For comparison purposes, cytokines of supernatants of unstimulated MAT-derived spheroids plotted are the same in Fig. 2C and Suppl. Fig. 2B. Spheroids were generated from n≥5 patients. Quantification is shown as mean ± SD. Statistical significance was calculated using unpaired student´s *t* test. (C) Experimental setup of the adipose transwell culture system. (D) Image of murine primary mature adipocytes cultured under a permeable transwell membrane (10x magnification) (E) Cytokine secretion profile of murine mature adipocytes under non-stimulated and stimulated conditions was measured using a multiplex immunoassay on transwell culture supernatants. *Transwell cultures were generated from n≥8 WT mice, pooled from 4 independent experiments. Quantification is shown as mean ± SD. Statistical significance was calculated using paired student´s t test.* *p < 0.05; **p < 0.01; ***p < 0.001; ****p < 0.0001. Illustration 2C was created with BioRender.com.

### Suppl. Fig. 3 - related to Fig. 3

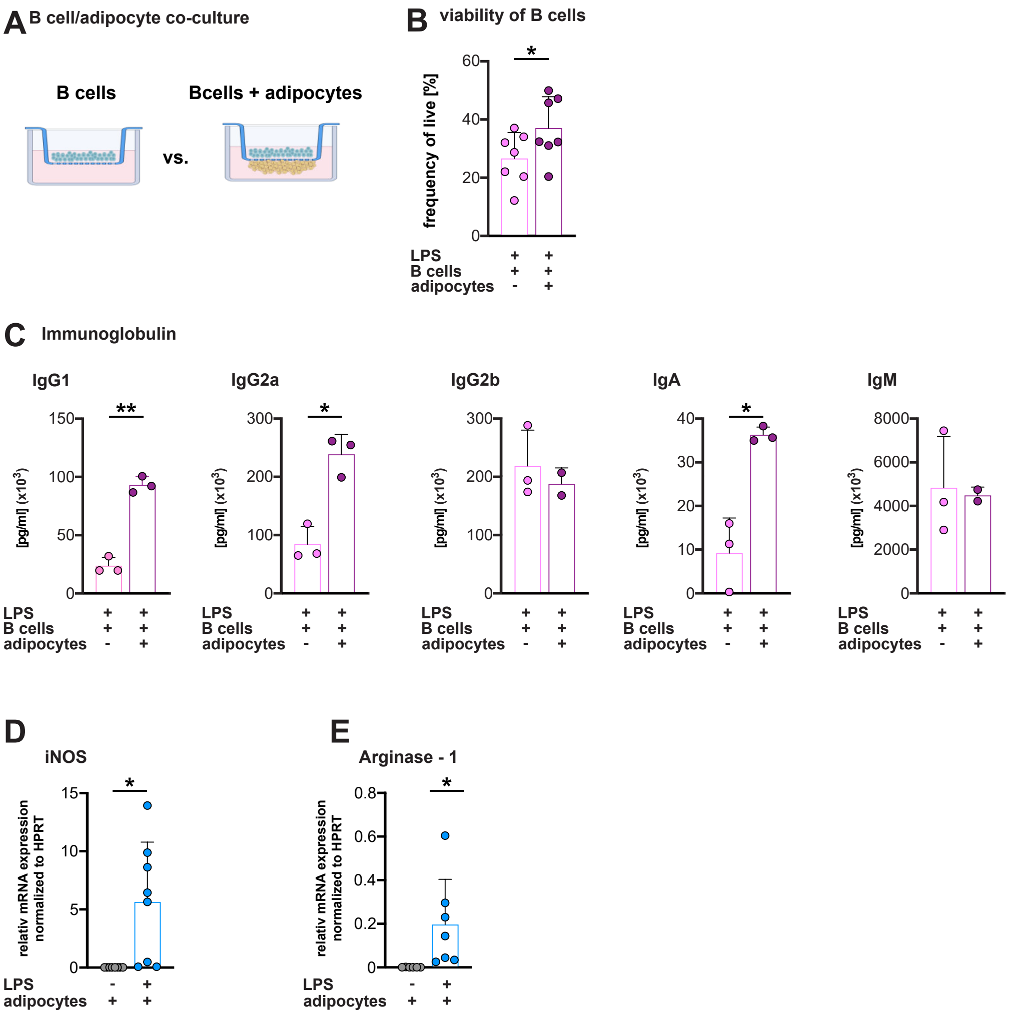


**Suppl. Fig. 3:**

(A) Schematic representation of the experimental design for B cell/adipocyte transwell co-cultures. (B) Bar graphs showing the frequency of viable stimulated B cells as determined by flow cytometry after culture either alone or in the presence of adipocytes. *Data is shown as mean ± SD. n≥7 mice pooled from 3-4 independent experiments.* (C) Concentrations of immunoglobulins released into the supernatants from mono- and co-cultures of stimulated B cells with adipocytes were measured using multiplex assays. *Data is shown as mean ± SD, n≥2 mice.*

(D+E) Relative mRNA expression levels of (E) iNOS and (F) Arginase-1 in BMDMs were measured after 24 hours of exposure to conditioned media as indicated. Results are shown as mean ± SD, n≥7 mice per condition pooled from 4 independent experiments.

*Statistical significance was calculated using paired student´s t test.* *p < 0.05; **p < 0.01; ***p < 0.001; ****p < 0.0001. Illustration 3A was created with BioRender.com.

### Suppl. Fig. 4 - related to Fig. 5

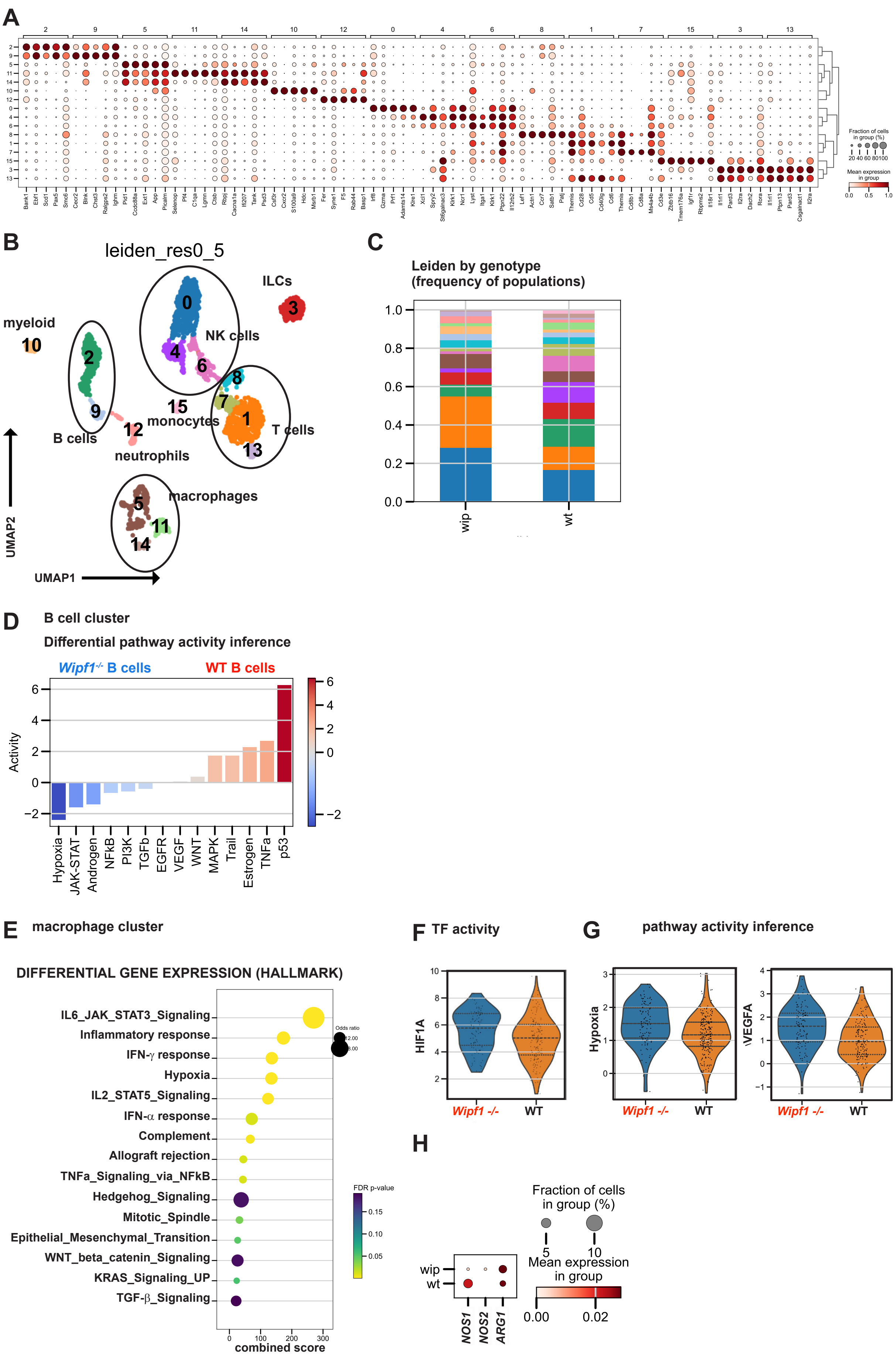

**Suppl. Fig. 4:**

(A) Dot plot showing the top five genes of each identified cell cluster most differentially expressed as identified by scanpy in single-cell RNA sequencing of CD45+ leukocytes isolated from murine WAT. Dot size indicates the proportion of cells expressing each gene within a cluster, and color intensity represents the average expression level. (B) UMAP-representation of immune-cell clusters identified using scRNAseq analysis of $CD45^{+}$ leukocytes isolated from WAT of WT and *Wipf1*$^{-/-}$ mice. (C) Frequency of leiden cluster distribution between the genotypes (D) Bar chart showing the distribution of inferred activity of indicated pathways in B cells between the genotypes. (E) Overrepresentation analysis of functional terms (Hallmark gene sets) in pooled macrophage clusters. Dot size reflects odds ratio, color shows FDR-adjusted p-value, x-axis displays combined overrepresentation score.(F) Violin plots showing the distribution of inferred transcription factor activity of *HIF1A* in macrophages between the genotypes. (G) Violin plots showing the distribution of inferred hypoxia and *VEGFA* pathway activity in B cells between the genotypes. (H) mRNA expression levels of indicated genes focusing on the macrophage cluster. Dot size indicates the proportion of cells expressing each gene within the group, while color intensity reflects the average expression level. n≥2 mice per genotype.

### Suppl. Fig. 5 - related to Fig. 5

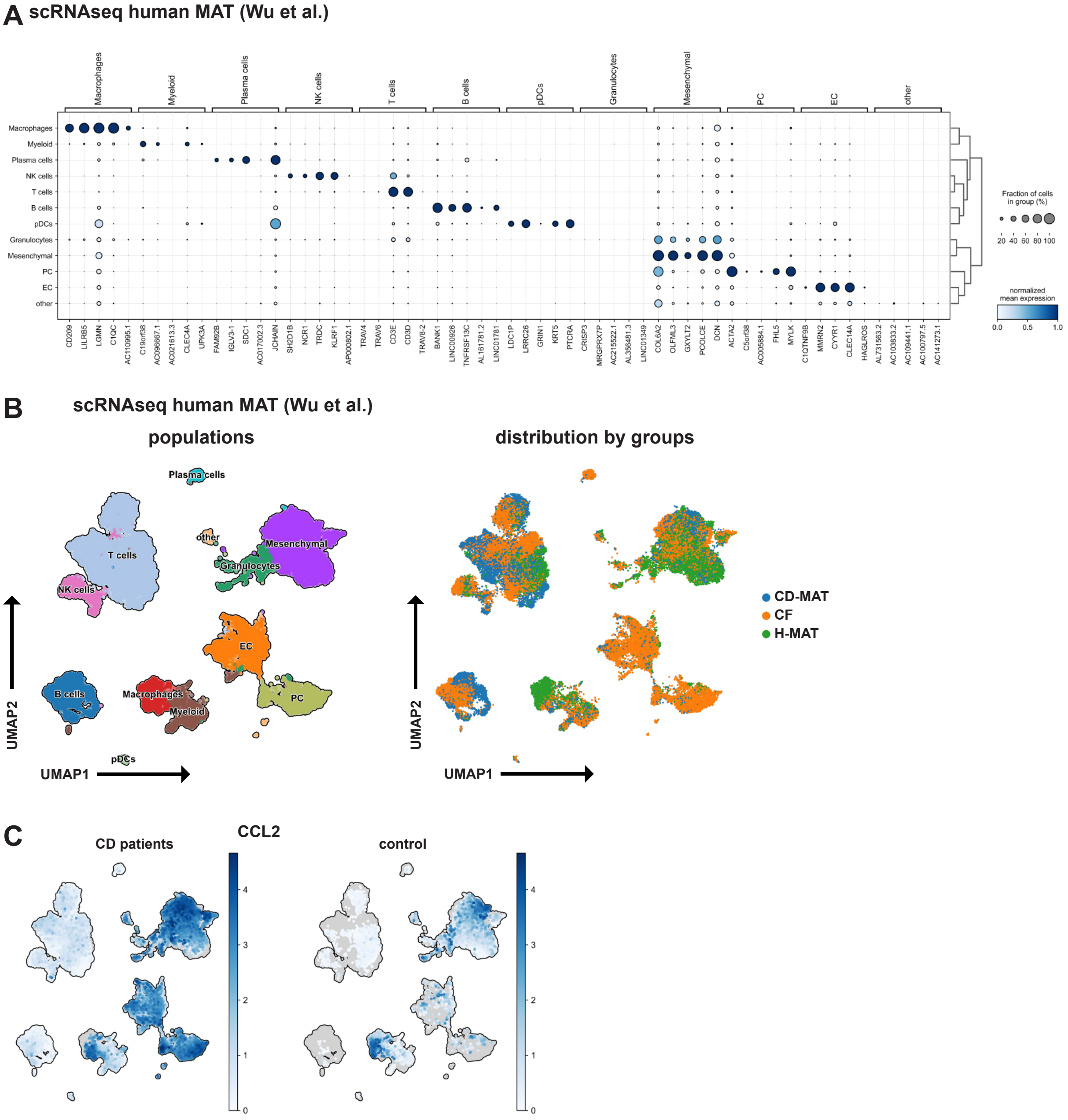


**Suppl. Fig. 5:**

(A)Dot plot showing the top five genes of each identified cell cluster most differentially expressed as identified by scanpy in single-cell RNA sequencing of single-cells isolated from HD-MAT, CD MAT and CrF (data extracted from (Wu et al., 2023)). Dot size indicates the proportion of cells expressing each gene within a cluster, and color intensity represents the average expression level. (B) UMAP-representation of cell clusters identified and the distribution by the different groups using scRNAseq analysis of single-cells isolated from HD-MAT, CD MAT and CrF. (C) UMAP-representation of expression levels of *CCL2* in CrF/MAT of CD patients compared to control MAT.

*Supplemental Table S1:*

**Patients` Characteristics - Samples used for immune cell phenotyping by flow cytometry**

| Patient ID | Age at resection | sex | diagnosis | year at first diagnosis | localization | sample | Medication at resection |
|---|---|---|---|---|---|---|---|
| H4 | 42 | m | Stricturing CD | 2002 | Ascending colon | MAT/CrF | Infliximab |
| H5 | 70 | f | Stricturing CD | 1991 | Terminal ileum | MAT/CrF | Glucocorticoide |
| H7 | 50 | f | Stricturing CD | 2021 | Transverse colon | MAT/CrF | Glucocorticoide |
| H9 | 31 | m | Stricturing CD | 2014 | Terminal ileum | MAT/CrF | Adalimumab |
| H13 | 24 | m | Stricturing CD | 2019 | Terminal ileum | MAT/CrF | Glucocorticoide |
| H15 | 54 | m | Stricturing CD | 1997 | Terminal ileum | MAT/CrF | Infliximab |
| H16 | 55 | f | Stricturing CD | 1982 | Terminal ileum | MAT/CrF | Ustekinumab |
| H19 | 71 | m | Stricturing CD | - | Terminal ileum | MAT/CrF | Ustekinumab |
| H20 | 29 | m | Stricturing CD | 2016 | Terminal ileum | MAT/CrF | Ustekinumab |

*Supplemental Table S2:*

**Patients Characteristics - Samples used for adipocyte-derived spheroids**

| Patient ID | Age at resection | sex | diagnosis | year at first diagnosis | localization | sample | Medication at resection |
|---|---|---|---|---|---|---|---|
| 25 | 62 | f | Stricturing CD | 2004 | Terminal ileum | MAT/CrF | - |
| 26 | 70 | f | Stricturing CD | - | Terminal ileum | MAT | - |
| 29 | 42 | m | Stricturing CD | 2002 | Ascending colon | MAT | Infliximab |
| 31 | 57 | f | Stricturing CD | 1994 | Terminal ileum | CrF | Steroids |
| 32 | 50 | f | Stricturing CD | 2021 | Transverse colon | MAT | Glucocorticoide |
| 34 | 31 | m | Stricturing CD | 2020 | Terminal ileum | MAT | Vedolizumab |
| 35 | 46 | m | Stricturing CD | 2001 | Sigmoid colon | CrF | Ustekinumab |
| 37 | 21 | m | Stricturing CD | 2019 | Terminal ileum | CrF | Glucocorticoide |
| 41 | 21 | m | Stricturing CD | 2008 | Terminal ileum | CrF | Ustekinumab |
| 46 | 28 | m | Stricturing CD | 2016 | Terminal ileum | MAT | Ustekinumab |
| 56 | 54 | m | Stricturing CD | 1997 | Terminal ileum | CrF | Infliximab |
| 60 | 55 | f | Stricturing CD | 1991 | Terminal ileum | CrF | Glucocorticoide |

***Supplemental Table S3***

***Antibodies and Dyes***

| **Antibody (Clone)** | **Target species** | **Fluorophore** | **Company** |
|---|---|---|---|
| BODIPY™ FLC16 | | | Thermo Fisher |
| Zombie Aqua™ Fixable Viability Dye | | | BioLegend |
| CD19 (HIB19) | human | APC-Cy7 | BioLegend |
| CD20 (2H7) | human | FITC | BioLegend |
| CD206 (MMR) | human | FITC | BioLegend |
| CD3 (HIT3a) | human | BV605 | BioLegend |
| CD3 (HIT3a) | human | BV785 | BioLegend |
| CD4 (SK3) | human | BV605 | BioLegend |
| CD45 (2D1) | human | PerCP-Cy5.5 | BioLegend |
| HLA DP DQ DR (TU39) | human | AF647 | BioLegend |
| HLA DP DQ DR (TU39) | human | BV421 | BD Biosciences |
| IgA (polyclonal) | human | AF555 | Southern biotech |
| Human TruStain FcX | human | | BioLegend |
| B220/CD45R (RA3-6B2) | mouse | BUV395 | BioLegend |
| B220/CD45R (RA3-6B2) | mouse | BV421 | BioLegend |
| CD11b (M1/70) | mouse/human | AF700 | BioLegend |
| CD163 (S15049I) | mouse | BV421 | BioLegend |
| CD19 (6D5) | mouse | BV785 | BioLegend |
| CD20 (sa275a11) | mouse | AF700 | BioLegend |
| CD206 (C068C2) | mouse | AF647 | BioLegend |
| CD3e (145-2C11) | mouse | PerCP-Cy5.5 | BioLegend |
| CD4 (GK1.5) | mouse | BV605 | BioLegend |
| CD45 (30-F11) | mouse | BUV395 | BioLegend |
| CD86 (GL-1) | mouse | PerCP-Cy5.5 | BioLegend |
| MHCII (M5/114.15.2) | mouse | APC-Cy7 | BioLegend |
| MHCII (M5/114.15.2) | mouse | AF488 | BioLegend |
| Purified CD16.2 (9E9) | mouse | | BioLegend |
| Purified CD16/32 (93) | mouse | | BioLegend |
| Ultra-LEAF™ Purified IFNγ | mouse | | BioLegend |
| Ultra-LEAF™ Purified IL-10 | mouse | | BioLegend |
| Ultra-LEAF™ Purified IL-6 | mouse | | BioLegend |
| Ultra-LEAF™ Purified MCP-1 | mouse | | BioLegend |
| Ultra-LEAF™ Purified TNFα | mouse | | BioLegend |

***Supplemental Table S4***

***Oligonucleotides***

Primers were synthesized at Eurofins Genomics (Ebersberg, Germany).

*murine qPCR Primer*

| **Enzymes for macrophage classification** | |
|---|---|
| iNOS_for | CTT TGC CAC GGA CGA GAC |
| iNOS_rev | TCA TTG TAC TCT GAG GGC TGA C |
| Arg-1_for | GGA ATC TGC ATG GGC AAC CTG TGT |
| Arg-1_rev | AGG GTC TAC GTC TCG CAA GCC |
| Control | |
| Hprt_for | AGCCTAAGATGAGCGCAAGT |
| Hprt_rev | TTACTAGGCAGATGGCCACA |
| **Hypoxia** | |
| HIFa_for | CCT GCA CTG AAT CAA GAG GTT GC |
| HIFa_rev | CCA TCA GAA GGA CTT GCT GGC T |
| NOX2_for | CCCTTTGGTACAGCCAGTGAAGAT |
| NOX2_rev | CAATCCCGGCTCCCACTAACATCA |
